# *Pseudomonas aeruginosa* biofilm-deficient mutants undergo parallel adaptation during chronic infection

**DOI:** 10.1101/2025.09.20.677542

**Authors:** Erin S. Gloag, Christopher W. Marshall, Nanami Kubota, Stacie E. Deaver, Brennan Deshotel, Vaughn S. Cooper, Daniel J. Wozniak

## Abstract

*Pseudomonas aeruginosa* readily adapts to infection by acquiring stable and heritable mutations. Previously, we discovered that the first adaptations in a porcine wound model were rugose small-colony variants (RSCVs) caused by mutations in the *wsp* operon. These mutants overproduce Pel and Psl biofilm exopolysaccharides that improve defense against host responses. To identify other mechanisms of host adaptation that lead to hyperbiofilm phenotypes, we created a mutant with an activated *wsp* pathway but unable to produce these exopolysaccharides (Δ*wspF*Δ*pelA*Δ*pslBCD*). Porcine wounds were infected with this mutant and biopsies were sampled at days 7, 14 and 35. Small colony variants were isolated from the wound, and whole genome sequencing revealed these variants had acquired mutations in genes in lipopolysaccharide and type IV pili biosynthesis, with *wzy* and *pilU* genes being most commonly targeted. *pilU* mutants were associated with a hyperbiofilm phenotype that outcompeted the parental strain, and *wzy* mutants were associated with a hyperbiofilm phenotype and increased tolerance to host antimicrobial products. We further identified that several variants had acquired large genome deletions that spanned up to 320 consecutive genes and other variants with high copy numbers of Pf6 filamentous phage. Together our results suggest that the hyperbiofilm phenotype is adaptive in chronic infections and that *P. aeruginosa* has redundant and diverse pathways to generate this phenotype.

## Introduction

*Pseudomonas aeruginosa* is an opportunistic pathogen, and evidence suggests that infections are established in compromised individuals through environmental reservoirs of the organism (1). Upon colonizing a host, *P. aeruginosa* often diversifies, acquiring beneficial adaptations that promote survival and persistence. An example of this is *P. aeruginosa* evolution and adaptation in response to colonizing the cystic fibrosis (CF) lung (2, 3). In the CF lung, the general paradigm is that *P. aeruginosa* evolves to become less virulent to promote chronic persistence (4) In line with this, mutations in specific genes or pathways are repeatedly targeted, including motility and surface attachment and cell wall or lipopolysaccharide synthesis (2, 3), suggesting that mutations in these genes are under selection during infection. Furthermore, mutations leading to increased biofilm formation, or a hyperbiofilm phenotype, are enriched in the CF lung (2, 3). In *P. aeruginosa*, one of the mechanisms controlling biofilm formation is the small signaling molecule c-di-GMP. The general paradigm is that low levels of c-di-GMP promote a planktonic lifestyle, while high levels of c-di-GMP promote a biofilm lifestyle (5) In *P. aeruginosa*, high c-di-GMP results in increased production of Pel and Psl polysaccharides, subsequently promoting increased biofilm formation (6). Furthermore, mutations leading to continued high c-di-GMP levels result in a rugose small-colony variant (RSCV) phenotype. These variants, due to the overproduction of Pel and Psl are associated with a hyperbiofilm phenotype (7) and increased tolerance to antimicrobials (8). In the CF lung, *P. aeruginosa* RSCVs of are frequently isolated that have mutations in the *wsp* system (2, 3).

The *wsp* system is a chemosensory system that senses membrane stress, and regulates c-di-GMP production upon surface attachment (6, 9). Mutations in *wspF, wspA* and *wspE* that result in continued activation of WspR, the diguanylate cyclase of the Wsp system, and subsequently the RSCV phenotype, are routinely identified in *P. aeruginosa* CF clinical isolates, indicating that these mutations result in adaptive phenotypes, and that the Wsp pathway experiences strong selective pressure during infection (2, 3). Furthermore, *P. aeruginosa* RSCVs have been isolated from a range of infections other than CF pulmonary infections, including ventilated associated pneumonia (10, 11), urinary tract infections (11, 12), catheter-associated urinary tract infections (13, 14), and canine otitis media (15). Hyperbiofilm variants, consistent with RSCVs, have also been isolated from osteomyelitis (16). Therefore, since RSCVs are considered to be adaptations to the biofilm lifestyle (7, 17), forming recalcitrant biofilms appears to be a common adaptation to chronic infections.

In our prior work we used a porcine full-thickness thermal injury model to identify beneficial mutations that promote *P. aeruginosa* persistence during chronic infection (18, 19). Using variant colony morphology as an indicator for adaptive variation, we identified RSCVs as the only variant colony phenotype to emerge across the 28 day infection (18). Whole genome sequencing revealed that mutations in the *wsp* chemosensory system, leading to the RSCV phenotype, were the first mutations to be selected in the infection(18). In the current study, we determined the genetic targets of *P. aeruginosa* adaptation to the wound in the absence of Pel and Psl exopolysaccharides to test alternative pathways of hyperbiofilm formation

## Results

### Exopolysaccharide independent small colony variants (SCVs) are selected during porcine chronic wound infections

We previously determined that mutations in the *wsp* operon, resulting in the RSCV phenotype, were among the earliest mutations to be selected during chronic infection (18). We hypothesized that the fitness of these variants is associated with the hyperbiofilm phenotype resulting from the overproduction of Psl and Pel exopolysaccharides. We therefore wanted to identify other pathways leading to hyperbiofilm phenotypes that experience selection during infection. To achieve this we assessed how a *wsp* mutant, with increased c-di-GMP but unable to produce Psl and Pel exopolysaccharides adapted to the infection environment. Using a porcine thermal injury chronic wound model (20), wounds were inoculated with MPAO1Δ*wspF*Δ*pelA*Δ*pslBCD* (Δ*wspFpelpsl*) and the infection burden monitored 7, 14 and 35 days post infection (dpi). Due to the absence of Pel and Psl polysaccharides, this mutant is deficient in biofilm formation, despite high intracellular c-di-GMP levels (21). This deficiency in biofilm formation resulted in an approximately 2-log reduction in bacterial burden in the wound (Fig 1A), compared to previous infections in this model using wild type *P. aeruginosa* (22).

**Figure 1:**
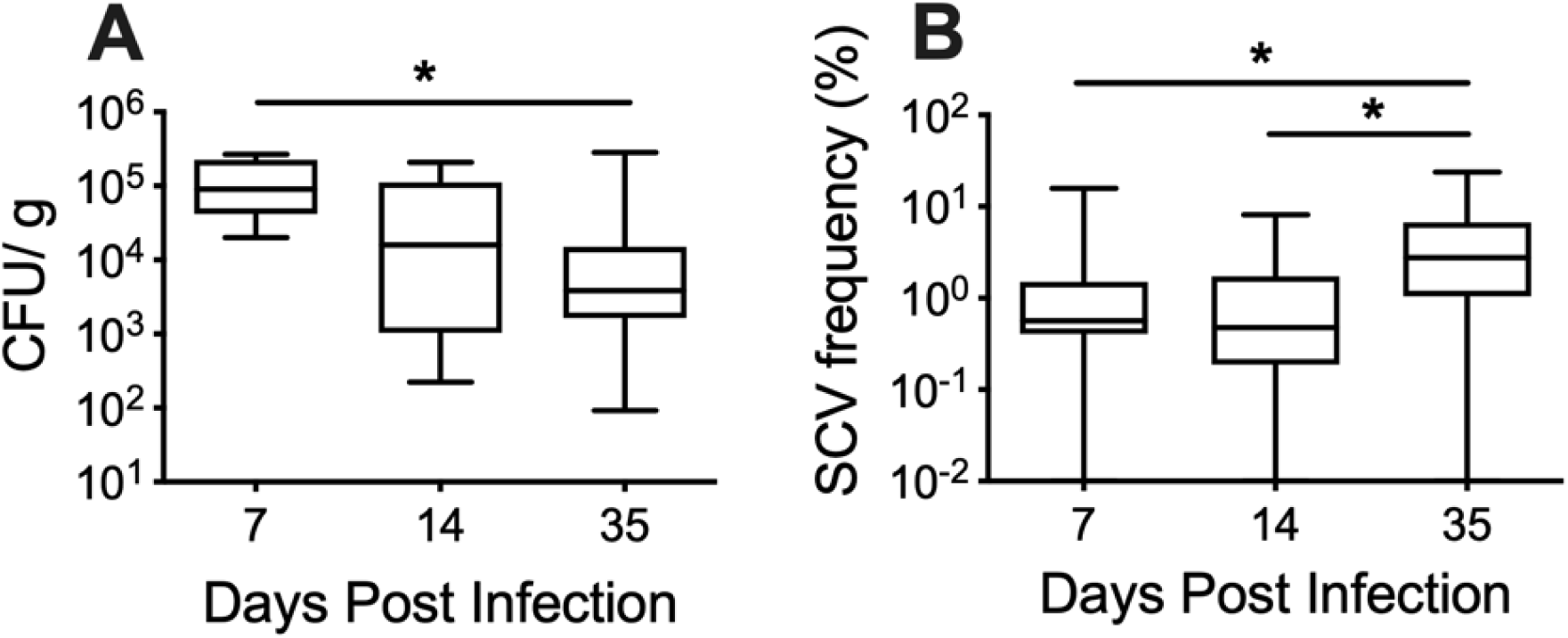
PAO1Δ*wspFpelpsl* burden in a chronic wound infection. Porcine wounds were inoculated with PAO1Δ*wspF*Δ*pelA*Δ*pslBCD* and biopsies were sampled from the wounds sampled at 7-, 14- and 35-days post infection. **(A)** Biopsies where homogenized and *P. aeruginosa* burden was enumerated CFU/ gram tissue. **(B)** Frequency of the SCV subpopulation screened from homogenized tissue at each time point, expressed as a percentage of the total *P. aeruginosa* population. Three biopsies were sampled from a total of four wounds per time point. N = 12, per time point. * *p*-value <0.05.

However, despite the lower bacterial burden, wounds still remained chronically colonized with Δ*wspFpelpsl* for the duration of the infection, with approximately 10^4^ CFU/g tissue recovered 35 dpi (Fig 1A).

To determine if populations founded by Δ*wspFpelpsl* adapted to the wound environment, variant colony morphology was used as an indicator for evolved variants that emerged during the infection. Homogenized biopsy samples were grown on *Pseudomonas* isolation agar and screened for colony variants. Small-colony variants (SCVs) were the only variant morphology observed. SCVs were isolated across all three time points, and the phenotype was stable across three passages on non-selective growth media, suggestive that the SCV phenotype was due to heritable mutation(s) (Fig 1B). The SCV frequency peaked at 35 dpi, at approximately 5% of the total *P. aeruginosa* population (Fig 1B). Upon closer inspection, two variant SCV morphologies were observed, a smooth SCV and wrinkled SCV, reminiscent of the RSCV phenotype (Fig 2A). Quantifying the frequency of each morphology as a percentage of the total SCV subpopulation revealed that the smooth SCVs predominated at all three time points. However, the wrinkled SCVs began to increase in frequency 35 dpi (Fig 2B).

**Figure 2:**
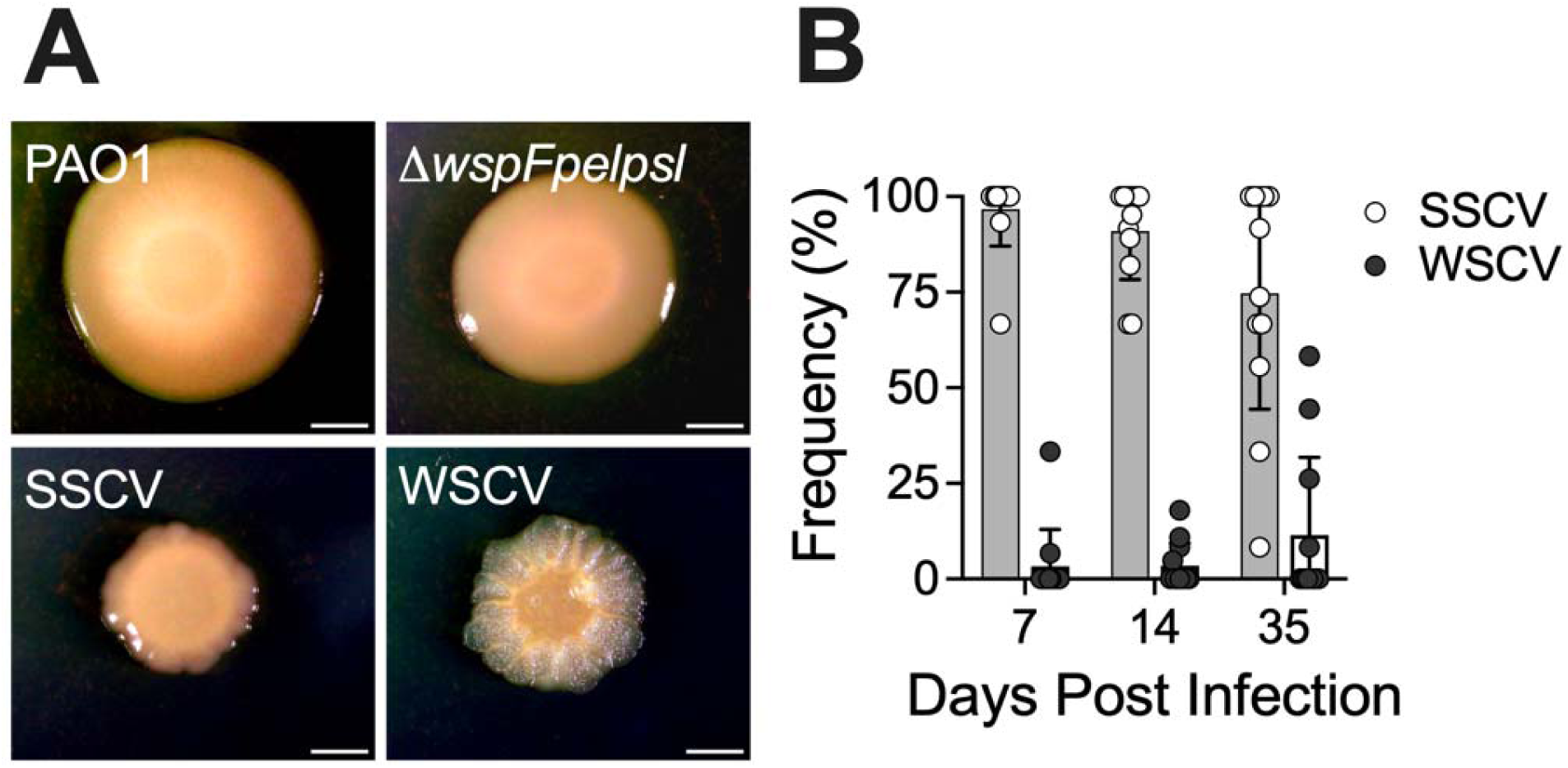
Colony morphology of SCVs isolated from porcine wounds. **(A)** Representative colony morphologies of SCVs isolated from porcine wounds, compared to wild type PAO1 and Δ*wspFpelpsl* ancestor (labelled). Representative smooth small-colony variant (SSCV) is SCV- 71, and representative wrinkled small-colony variant (WSCV) is SCV-51 (Table 2). Scale bar indicates 2 mm. **(B)** Frequency of the two variant colony morphologies, SSCV and WSCV, expressed as a percentage of the total SCV subpopulation at each time point.

To determine if the SCV subpopulation experienced positive selection in the wound, selective coefficients were determined as a measure of relative fitness (equation 1) using a range of potential starting SCV frequencies (Table 1). This revealed that across all three time points, SCVs likely had a selection rate >0.1 (i.e. 10% fitness advantage per generation) for all potential frequencies, indicating that SCVs experience strong positive selection in the wound.

**Table 1:** Selection coefficient (*s*) of SCVs from porcine wounds, inferred from potential starting frequencies.

| Starting frequency<br>(SCV: Ancestor) | s (mean +/- SD) days post infection |  |  |
| --- | --- | --- | --- |
|  | 7 | 14 | 35 |
| 1:10<sup>5</sup> | 0.96 +/- 0.15 | 0.50 +/- 0.14 | 0.26 +/- 0.05 |
| 1:10<sup>6</sup> | 1.28 +/- 0.16 | 0.65 +/- 0.10 | 0.31 +/- 0.03 |
| 1:10<sup>7</sup> | 1.60 +/- 0.19 | 0.81 +/- 0.09 | 0.36 +/- 0.03 |
| 1:10<sup>8</sup> | 1.92 +/- 0.23 | 0.96 +/- 0.09 | 0.41 +/- 0.05 |

### Exopolysaccharide independent SCVs acquire mutations in LPS O-antigen and type IV pili

To determine the mutation(s) responsible for the SCV phenotype, whole genome sequencing was performed on 39 randomly selected SCVs (Table 2). This revealed that most putative driver mutations responsible for the SCV phenotype occurred in genes involved in either the type IV pili (T4P) or lipopolysaccharide (LPS) biosynthesis pathways (Fig 3). Furthermore, mutations in genes in the T4P pathway became more prevalent at the later time points (Fig 3). Specifically, *pilU* (Fig 4A) and *wzy* (Fig 4B) were the most frequently mutated genes belonging to the T4P and LPS pathways, respectively. PilU is an ATPase that powers retraction of the pilus (23), while Wzy is the polymerase for the synthesis of LPS O-antigen (24). Interestingly, *pilU* mutations were associated with the wrinkled SCV phenotype (Fig 2A). Consistent with this, a matte colony morphology has previously been observed for *pilU* mutants (25, 26). The mutational parallelism in these two biosynthetic operons (Fig 5), observed across *P. aeruginosa* isolates, wounds and time points is a strong indicator that T4P and LPS are under selection in this strain in the chronic wound environment.

**Table 2:**
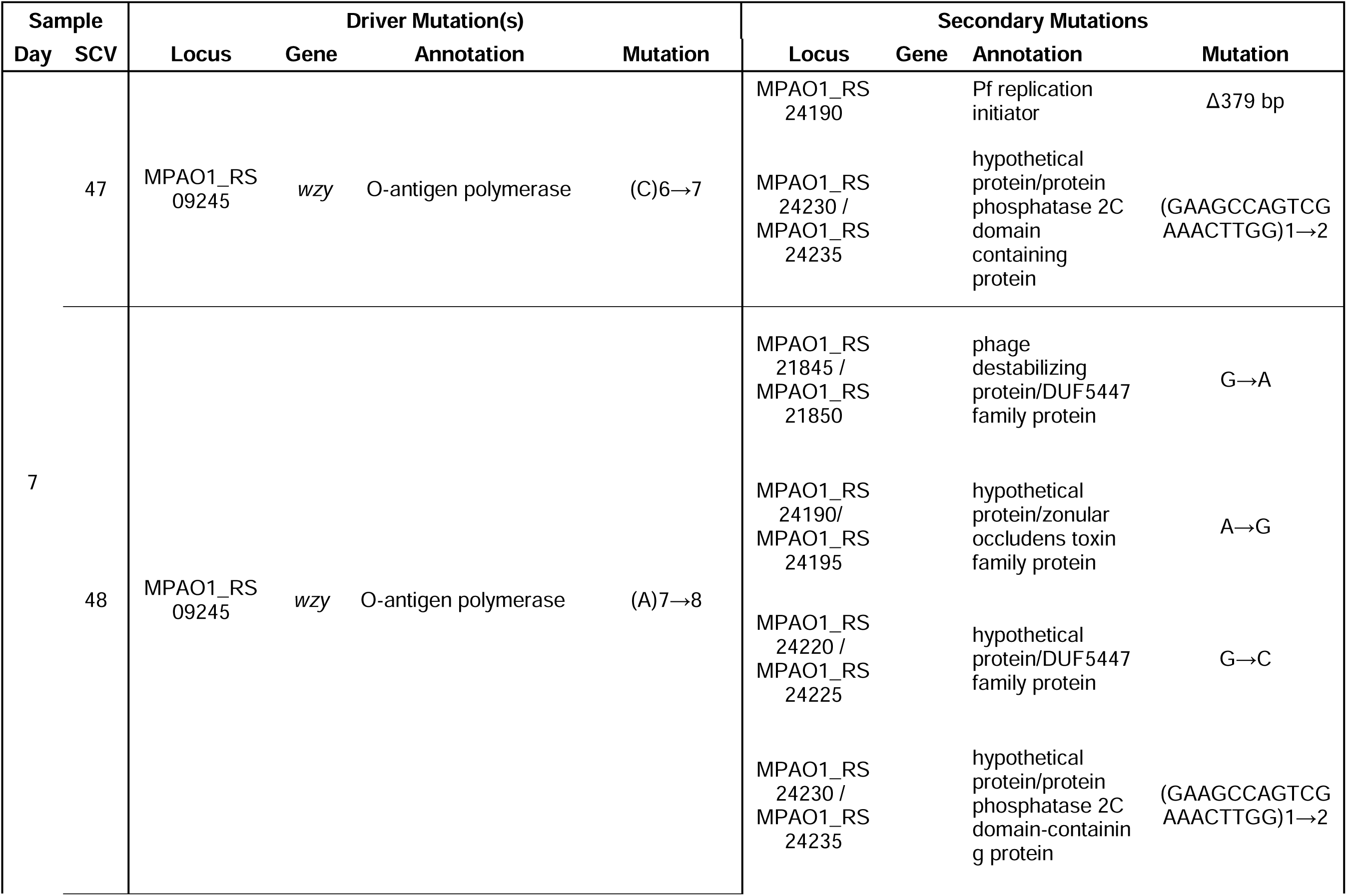
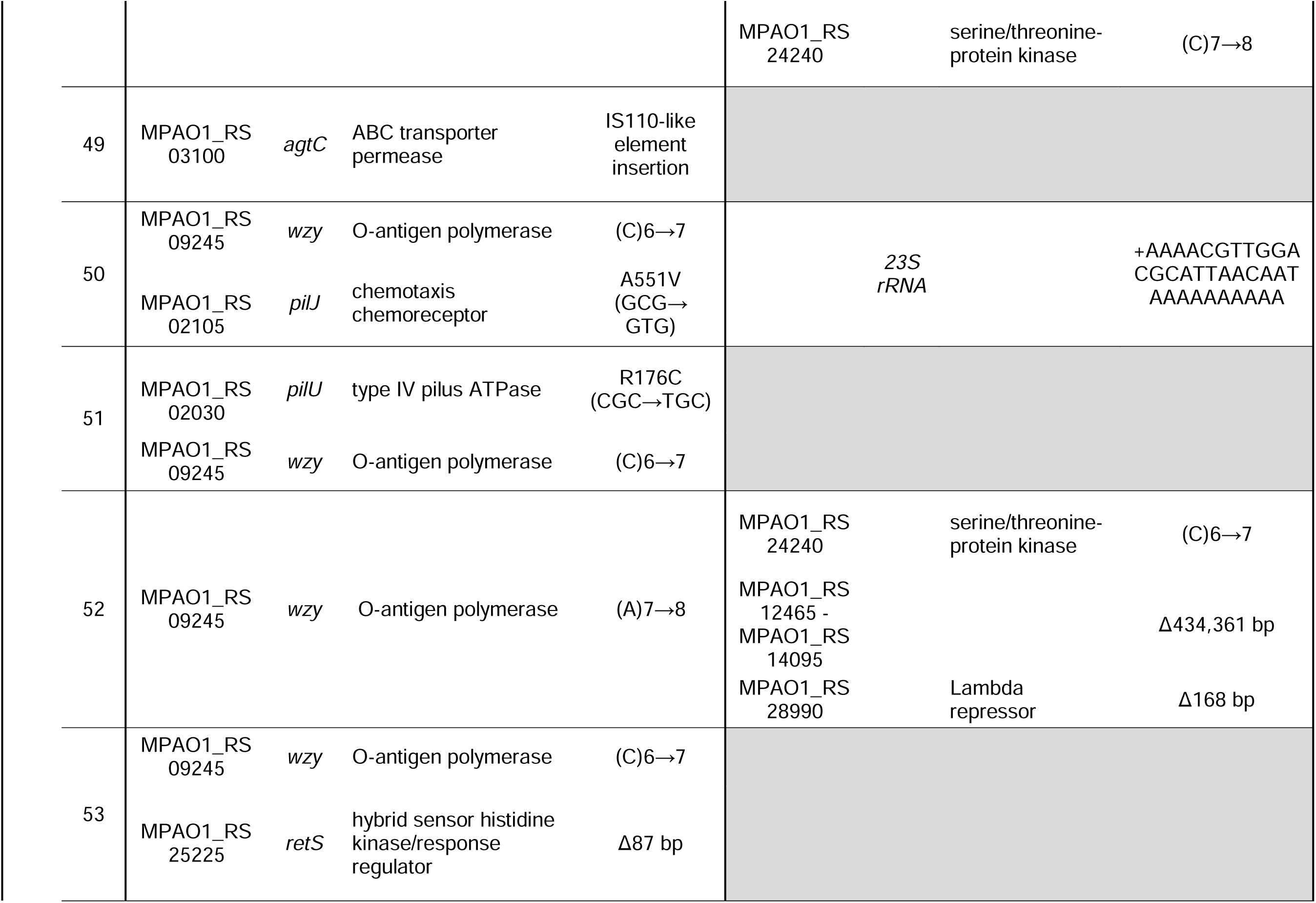
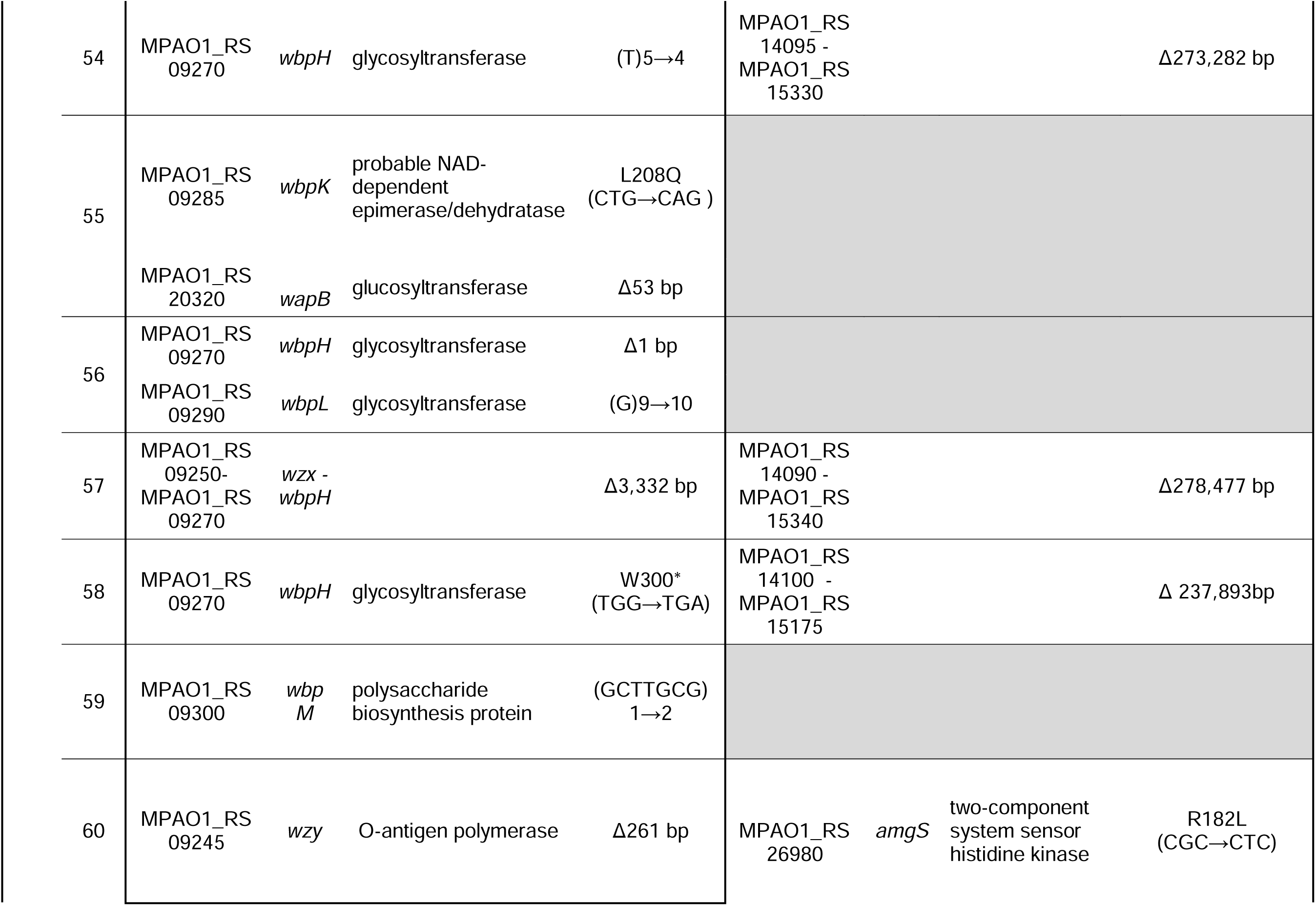
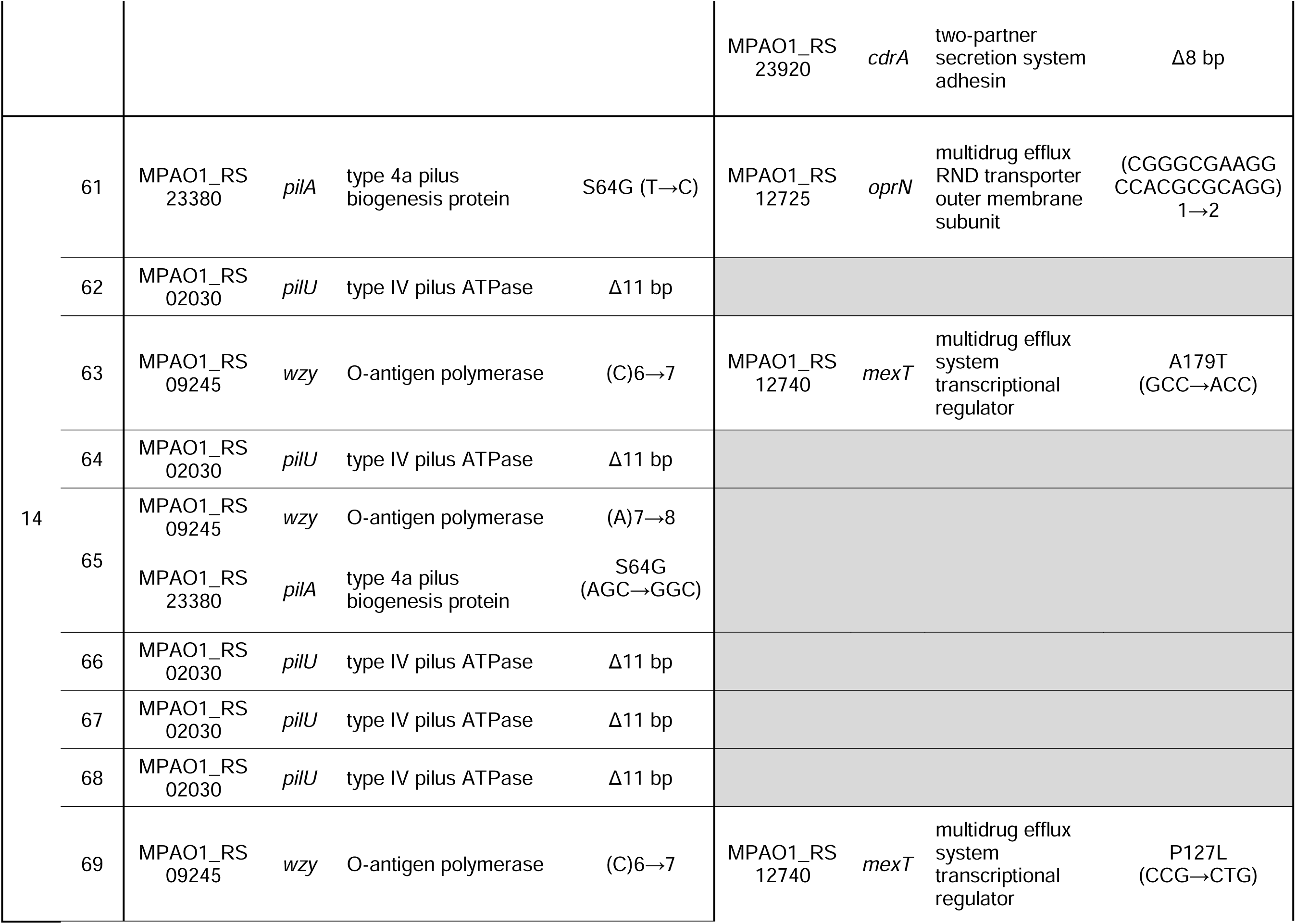
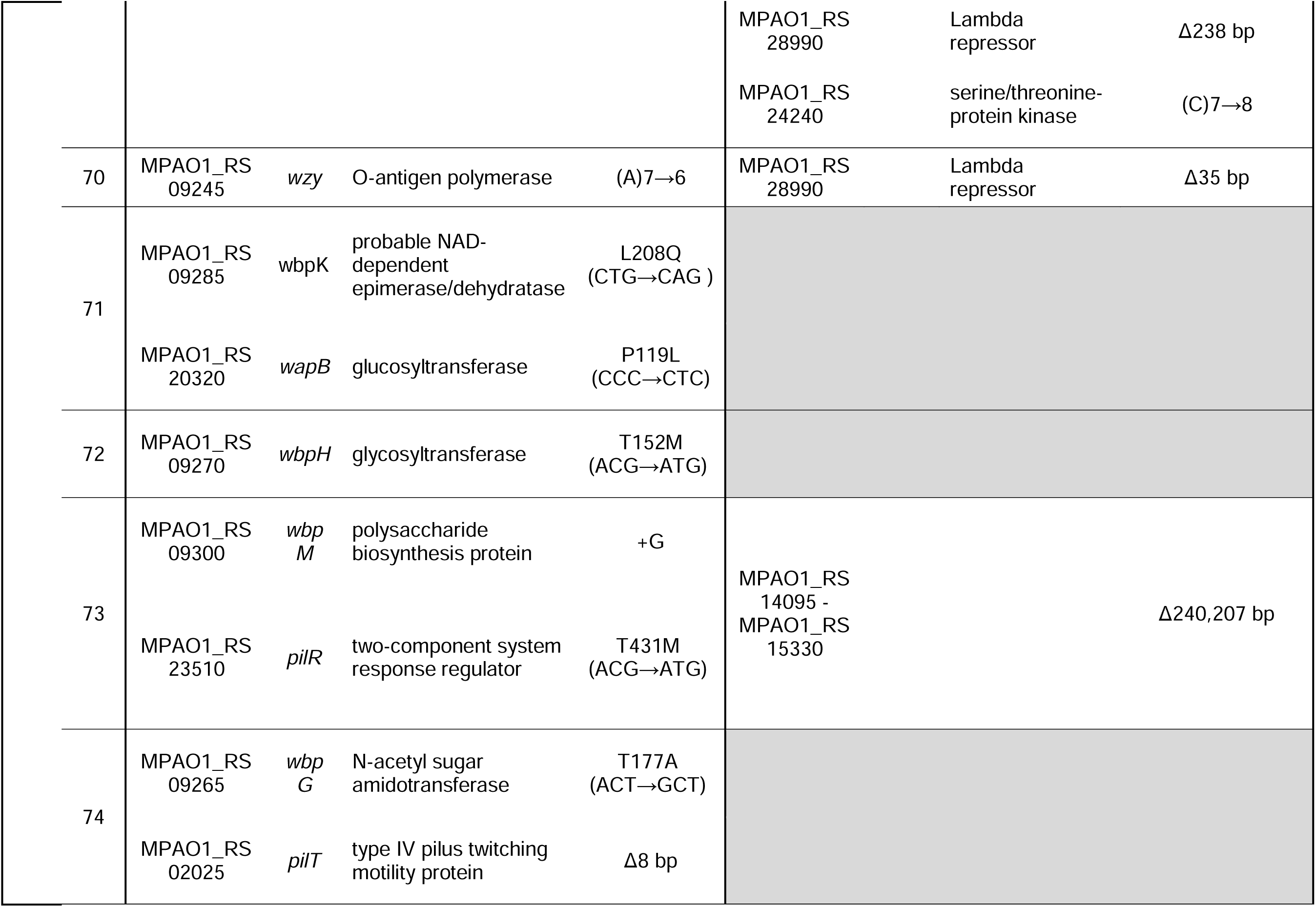
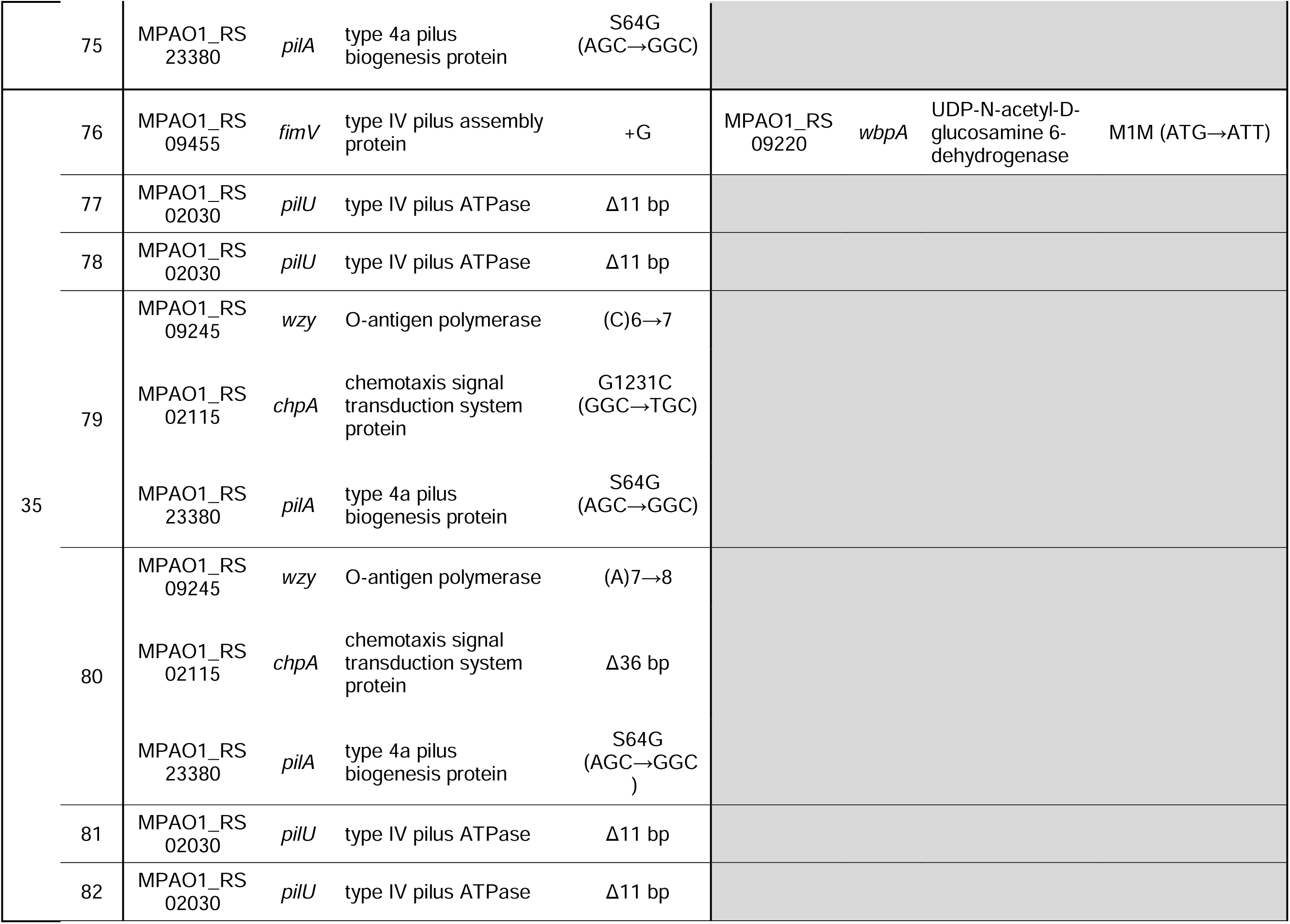
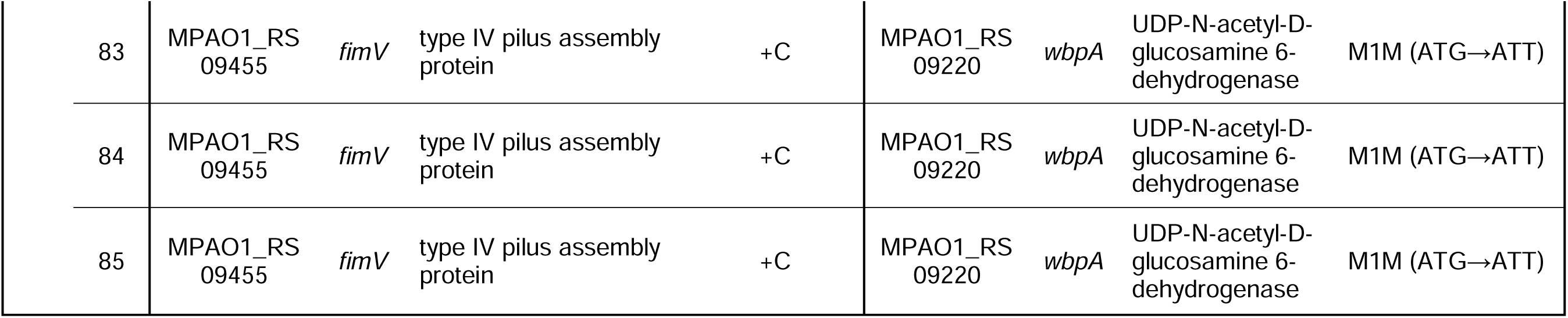
Mutations identified in SCVs.

**Figure 3:**
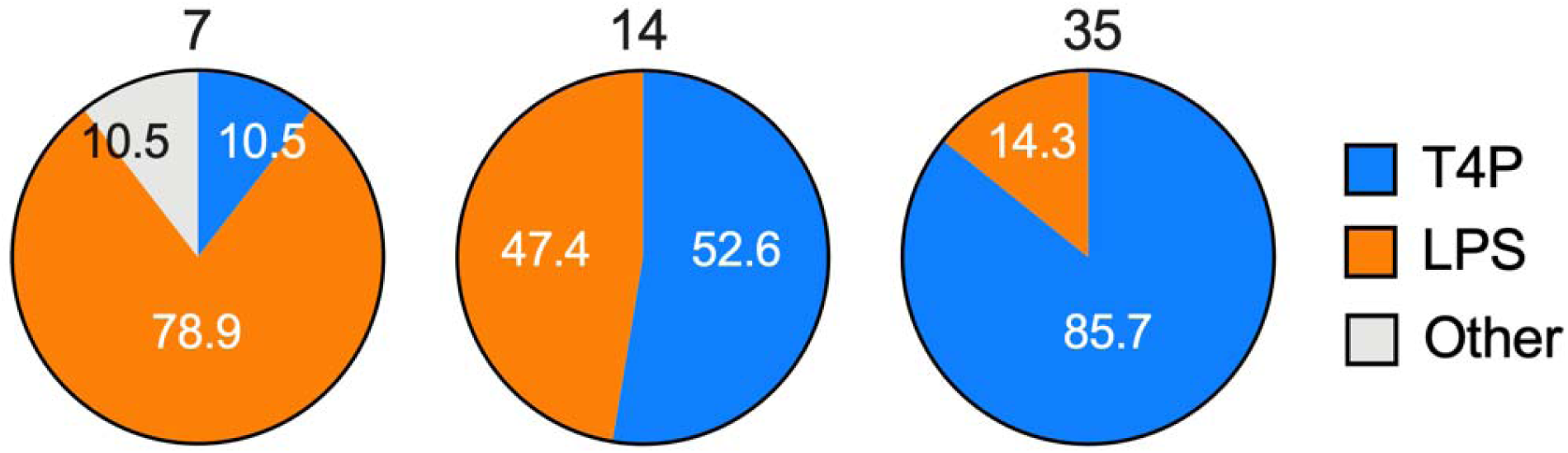
Classification of putative SCV mutations. 39 representative SCVs were sequenced. Mutations predicted to be responsible for variant colony morphologies could be categorized as either mutations in the T4P (blue) or LPS (orange) biosynthetic pathways, or other (grey). The number of mutations in either pathway is expressed as a percentage (labelled) of the total identified SCV mutations at each time point (labelled). Individual mutations are listed in Table 2.

**Figure 4:**
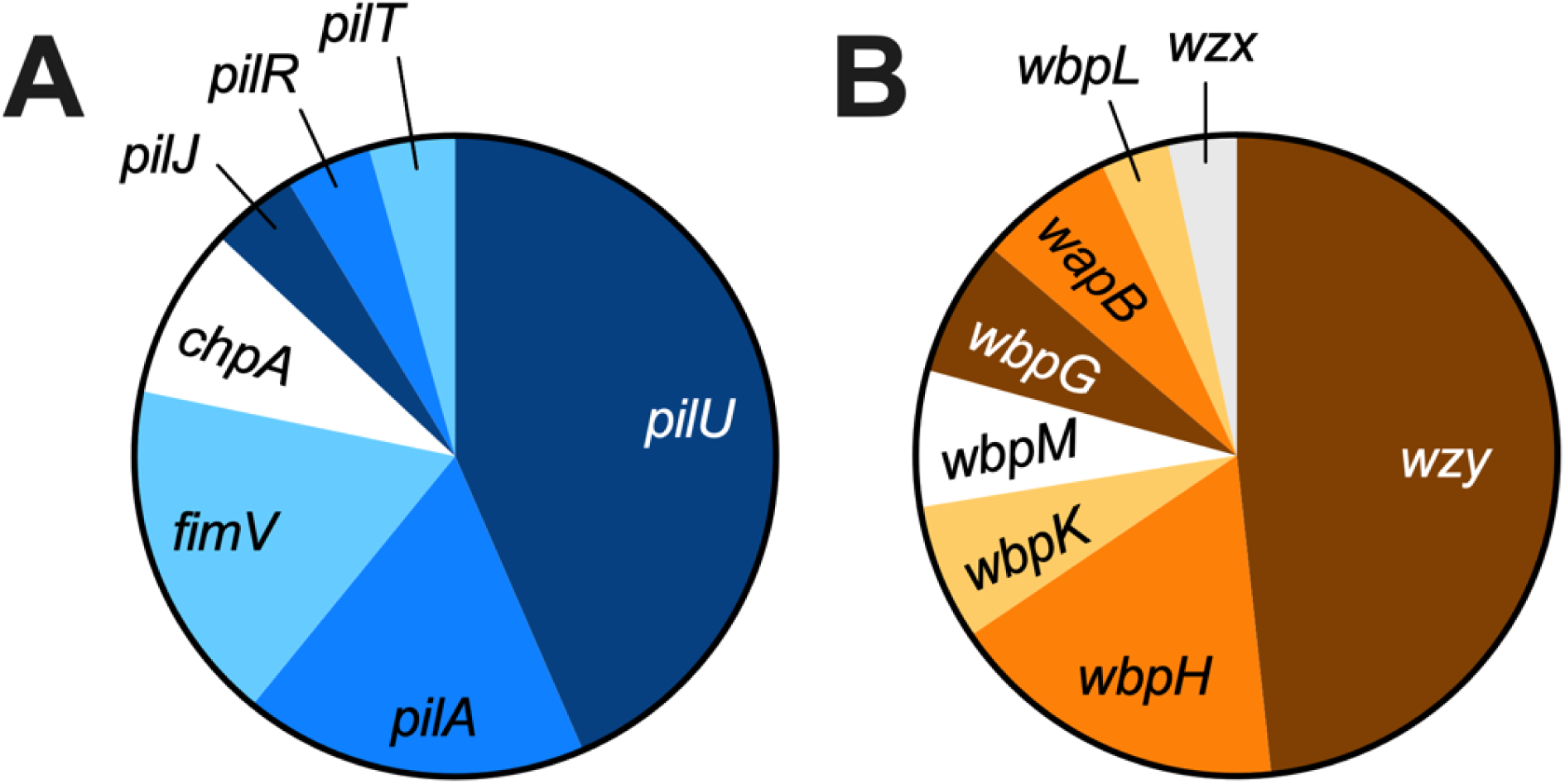
Frequency of genes with nonsynonymous SCV mutations. Frequency of driver mutations in genes (labelled) in either the **(A)** T4P or **(B)** LPS biosynthetic pathways, expressed as a percentage of the total number of SCV mutations, combined across all three time points. Individual mutations are listed in Table 2. Percentage of each gene is as follows **(A)** *pilU* 43.5; *pilA* 17.4; *fimV* 17.4; *chpA* 8.7; *pilJ* 4.3; *pilR* 4.3; *pilT* 4.3; **(B)** *wzy* 48.3; *wbpH* 17.2; *wbpK* 6.9; *wbpM* 6.9; *wbpG* 6.9; *wapB* 6.9; *wbpL* 3.4; *wzx* 3.4.

**Figure 5:**
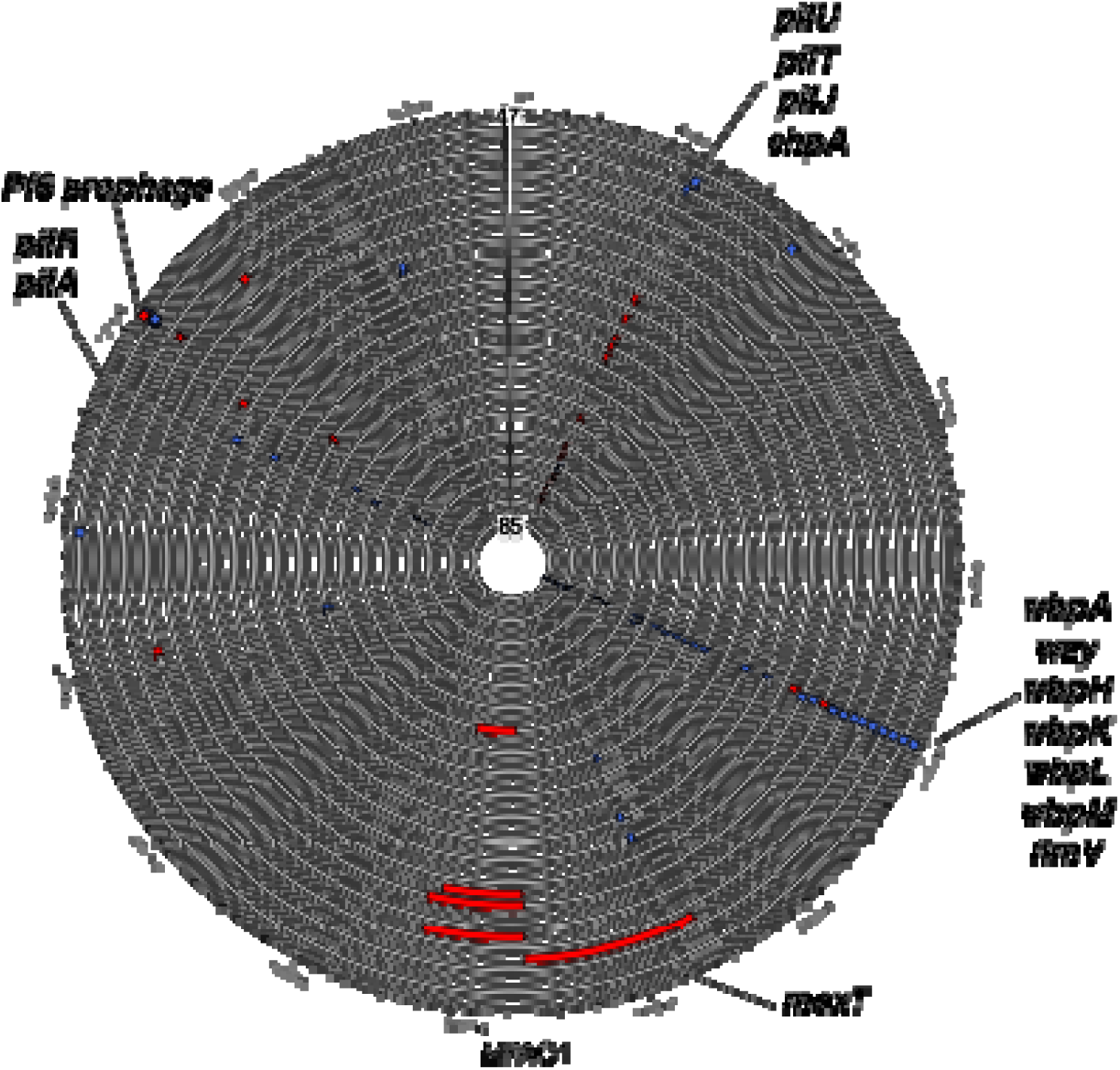
Parallel evolution across PAO1Δ*wspFpelpsl* isolates. Circular genome map of each sequenced SCV, represented by a double black line, aligned to the MPAO1 genome. For simplicity every second SCV is labelled. Mutations listed in Table 2 are represented, red are deletions, blue are small insertions or SNPs. Labelled are the location of specific genes. Large deletions for SCVs 52, 54, 57, 58 and 73 begin at bp locations 2,637,522, 3,072,354; 3,069,871; 3,072,771; 3,069,919 on the MPAO1 genome, respectively.

Sequencing of the SCVs revealed that many single isolates had acquired multiple mutations (Table 2). To determine the phenotypes and fitness effects associated with *pilU* and *wzy* mutations, and the contribution of the Δ*wspFpelpsl* background, the most common mutation in each gene was recreated in MPAO1 (referred subsequently here as PAO1), Δ*pelA*Δ*pslBCD* (Δ*pelpsl*), Δ*wspF,* and Δ*wspFpelpsl* backgrounds. The most common mutation in *pilU* was an 11bp deletion of bases 560 – 570 (*pilU*_Δ560-570_) (Table 2). In the PAO1 background, the *pilU*_Δ560-570_ mutant had a loss of twitching motility, comparable to that of a complete *pilU* gene deletion mutant (Δ*pilU*) (Fig S1), consistent with previous observations of *pilU* mutants (25, 26).

Furthermore, introduction of wild type *pilU in trans* restored twitching motility of both *pilU*_Δ560-570_ and Δ*pilU* mutants to that of wild type, indicating that *pilU*_Δ560-570_ is a loss-of-function mutation (Fig S1). The most common mutation in *wzy* was a single bp insertion at position 620 (*wzy*_620insC_) (Table 2). This resulted in a loss of O-antigen, or B-band LPS, production (Fig S2). Introduction of wild type *wzy in trans* restored O-antigen production to that of PAO1 (Fig S2), indicating that *wzy*_620insC_ is a loss-of-function mutation.

We initially hypothesized that *P. aeruginosa* adaptation to the wound is targeted towards mutations leading to hyperbiofilm phenotypes. To therefore determine if the *pilU*_Δ560-570_ mutation was associated with changes to biofilm formation, biofilms were grown for 24h in a 96-well plate, and biofilm biomass quantified by crystal violet. In all backgrounds, except for Δ*wspF*, the *pilU*_Δ560-570_ mutation was associated with a significant increase in biofilm formation, compared to the parental strain (Fig 6A). Quantification of biofilm bacteria by colony forming units revealed that in the Δ*wspFpelpsl* background, the *pilU*_Δ560-570_ mutation did not result in a significant increase of cells in the biofilm (Fig S3). We therefore predict that the increased biomass is attributed to hyper piliation of the cells due to a lack of a functional PilU (23), contributing a proteinaceous EPS. To determine if the *pilU*_Δ560-570_ mutation was associated with changes in fitness, the mutant was competed against the parent, which was tagged with *lacZ*, in a biofilm bead assay (27). Importantly, the *lacZ* tag did not confer any changes in fitness to the parent strain (Fig S4). The selection rate of the mutant was determined at 24h and 48h, according to equation 1. Across both time points and in all backgrounds, the *pilU*_Δ560-570_ mutation was associated with increased fitness, as indicated by *r* > 0.1, compared to the parent (Fig 6B). Interestingly, the highest fitness of the *pilU*_Δ560-570_ mutation was observed in the Δ*wspFpelpsl* background after 24h (Fig 6B).

**Figure 6:**
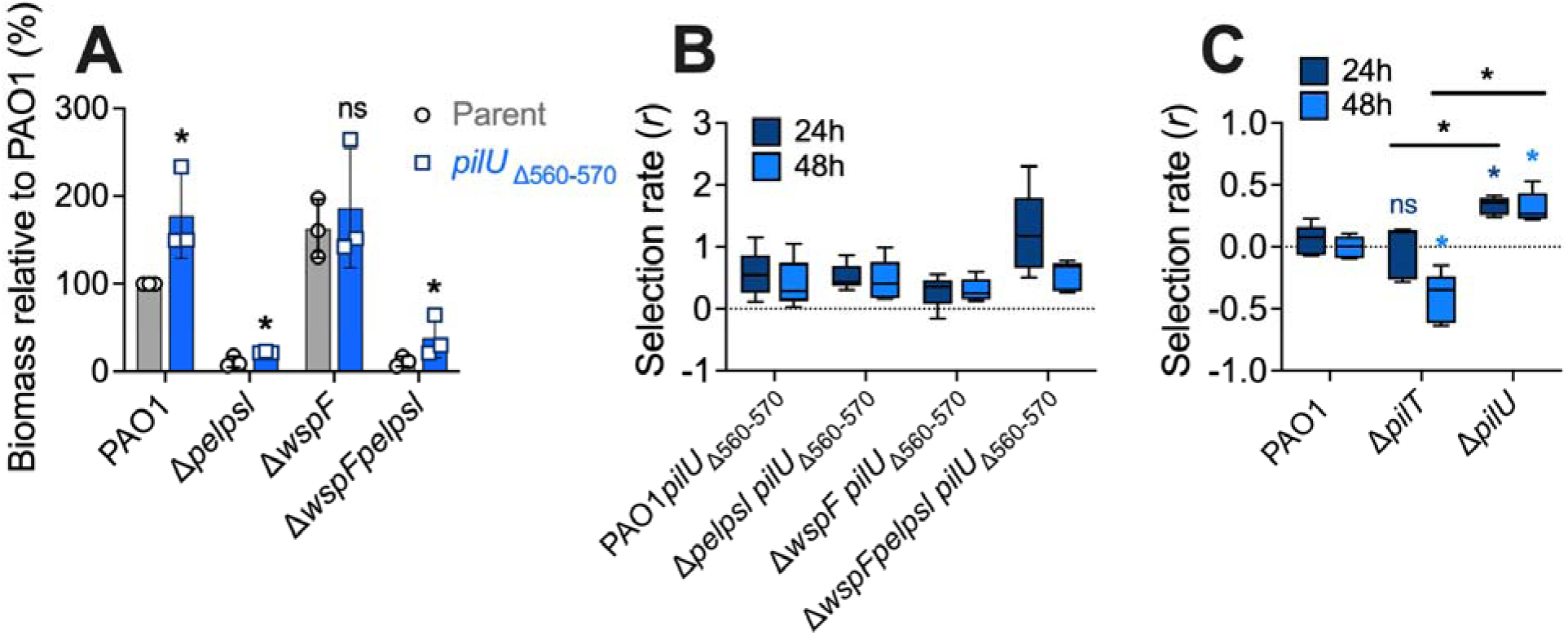
Mutations in *pilU* promote increased biofilm formation and fitness. **(A)** Biofilms were grown in a 96-well plate for 24h in Jensen’s media. Biofilm biomass was quantified by crystal violet staining. Biomass expressed as a percentage relative to PAO1. Data presented as mean ± SD. Individual data points indicate biological replicates, which are the average of four technical replicates. N = 3. Significance was determined using two-tailed unpaired t-test, between the parent and mutant. * *p*-value <0.05, ns indicates no significance, compared to the parent background. Selection rate (*r*) of **(B)** *pilU*_Δ560-570_ mutants or **(C)** complete *pilT* and *pilU* gene deletions, competed pairwise against the parent in a biofilm for 24 and 48h (dark and light blue, respectively). N = 5. **(C)** Significance determined using a two-way ANOVA with a Tukey’s multiple comparison posthoc test. * *p*-value <0.05, ns indicates no significance. Colored *, comparison to PAO1 competition at either 24 or 48h. Black *, comparison indicated on the graph.

*P. aeruginosa* encodes a second ATPase, *pilT*, that also powers the retraction of T4P (28). Both *pilT* and *pilU* mutants have a hyper piliation phenotype (23). To therefore determine if the increased fitness of the *pilU*_Δ560-570_ mutation was due to a general hyper piliation phenotype, or specific to a *pilU* mutation, complete gene deletions of *pilT* and *pilU* were constructed in PAO1, and competed pairwise against PAO1 tagged with *lacZ* in the biofilm bead assay (27). The selection rate of the mutant was determined at 24h and 48h, according to equation 1. After 24h, there was no change in fitness of Δ*pilT*, however after 48h there was a significant decrease in fitness, when compared to the parent competition control (Fig 6C). In contrast, Δ*pilU* had significantly increased fitness at both 24 and 48h, compared to both the parent control and Δ*pilT* (Fig 6C), like what was observed for the *pilU*_Δ560-570_ mutation (Fig 6B). Together this indicates that the increased fitness associated with the *pilU*_Δ560-570_ mutation is specific to *pilU*, and not due to the general hyper piliation phenotype.

We next determined phenotypes associated with the *wzy*_620insC_ mutation. To determine if the mutation was associated with changes to biofilm formation, biofilms were grown for 24h in a 96- well plate, and biofilm biomass quantified by crystal violet. This revealed that in Δ*wspFpelpsl* the *wzy*_620insC_ mutation resulted in a significant increase in biofilm formation, compared to the parental strain (Fig 7A). In PAO1 and Δ*pelpsl*, the *wzy*_620insC_ mutation was associated with increased biofilm formation, however this was not statistically significant. No difference in biofilm formation was observed for the *wzy*_620insC_ mutation in the Δ*wspF* background (Fig 7A).

**Figure 7:**
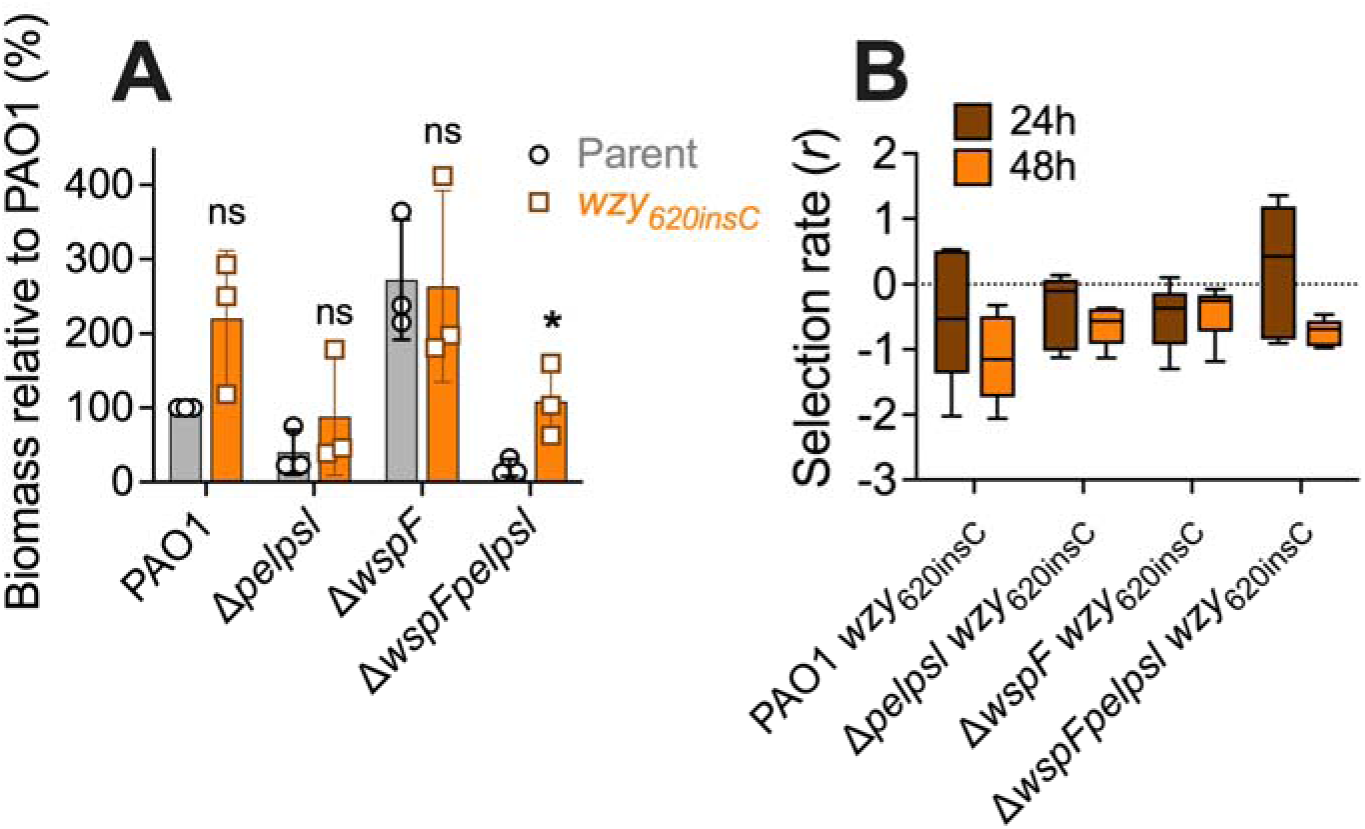
Mutations in *wzy* promote increased biofilm formation, but not increased fitness. **(A)** Biofilms were grown in a 96-well plate for 24h in Jensen’s media. Biofilm biomass was quantified by crystal violet staining. Biomass expressed as a percentage relative to PAO1. Data presented as mean ± SD. Individual data points indicate biological replicates, which are the average of four technical replicates. N = 3. Significance was determined using two-tailed unpaired t-test, between the parent and mutant. * *p*-value <0.05, ns indicates no significance, compared to the parent background. **(B)** Selection rate (*r*) of *wzy*_620insC_ mutants, competed pairwise against the parent in a biofilm for 24 and 48h (dark and light orange, respectively). N = 5.

Enumeration of biofilm bacteria revealed that in the Δ*wspFpelpsl* background, the *wzy*_620insC_ mutant resulted in a significant increase in cells (Fig S3), accounting for the increased biofilm phenotype in this strain background (Fig 7A). To determine if the *wzy*_620insC_ mutation was associated with changes in fitness, the mutant was competed against the *lacZ* tagged parent, in the biofilm bead assay (27). In all strain backgrounds, at both 24 and 48h, the *wzy*_620insC_ mutation was associated with decreased fitness, indicating that the mutant was outcompeted by the parent in this assay (Fig 7B). This is despite the apparent increased biofilm phenotype of *wzy*_620insC_ mutants (Fig 7A). We predict that a reduced growth rate of *wzy*_620insC_ mutants contributed to the reduced fitness observed when competed against the parental strain.

To identify potential fitness advantages of the *wzy*_620insC_ mutation in the wound-like environment, we tested bacterial survival in response to host antimicrobial products. PAO1, Δ*wspFpelpsl,* and Δ*wspFpelpsl wzy*_620insC_ were grown to mid-log and treated with either PBS (untreated control), H_2_O_2_, or serum for 1h, and bacterial survival was quantified (Fig 8). PAO1 and Δ*wspFpelpsl* had equivalent levels of survival in both the H_2_O_2_ and serum treatments. However, in the Δ*wspFpelpsl* background, the *wzy*_620insC_ mutation resulted in increased survival to H_2_O_2_ and serum, although this was only statistically significant for serum (Fig 8). This suggests that the fitness advantage associated with the *wzy*_620insC_ mutation may be due to increased survival to host antimicrobial products, including reactive oxygen species and those present in serum such as complement. We did not observe any increased survival associated with the *pilU*_Δ560-570_ mutation (Fig S5).

**Figure 8:**
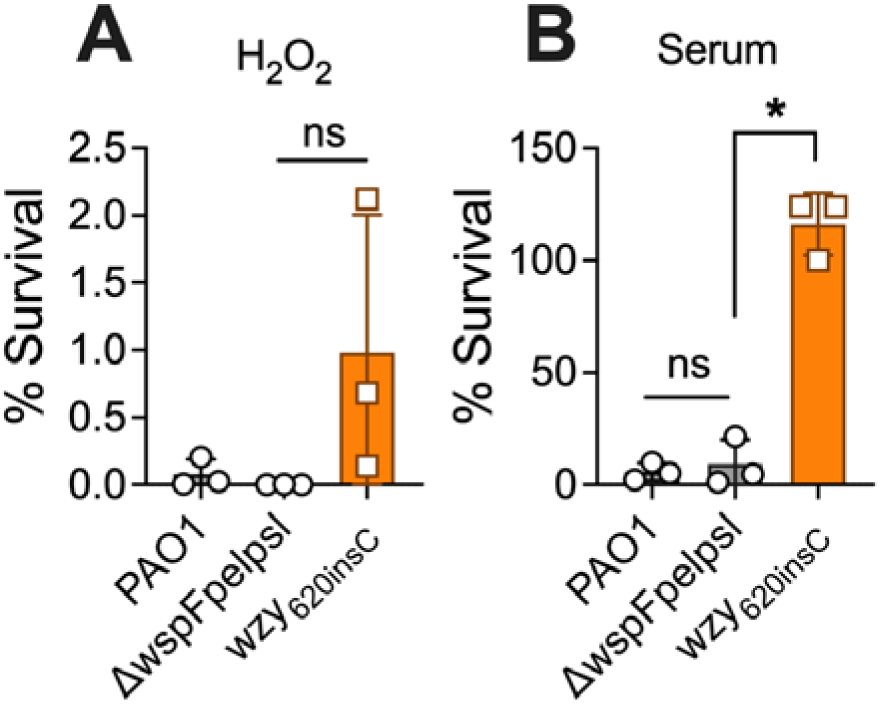
***wzy*_620insC_ mutation is protective against host antimicrobial products.** PAO1, Δ*wspFpelpsl* and Δ*wspFpelpsl wzy*_620insC_ was treated with either **(A)** 2.5% H_2_O_2_ or **(B)** 100% serum for 1h. Bacterial viability was enumerated by CFU/mL and expressed as percent survival relative to PBS untreated control. N = 3. Data presented as mean ± SD. Individual data points indicate biological replicates, which are the average of three technical replicates. * *p*-value <0.05, ns indicates no significance. Comparisons are indicated on the graph.

### Large genomic deletions associated with a subset of SCVs

In addition to the driver SCV mutations described above, we also identified that 5 of the SCVs had acquired unique deletions of large segments (>200kb) of DNA, with up to 320 consecutive genes deleted (Table 2, Fig 5). SCVs 54, 57, 58 and 73 had genomic deletions spanning the same region, while SCV 52 had a deletion in the adjacent genomic region (Fig 5), again highlighting the level of mutational parallelism experienced during infection.

Phenotypic analysis of these SCVs revealed that SCVs 57 and 73 had increased biofilm formation relative to the Δ*wspFpelpsl* parent (Fig S6A), however when competed pairwise with Δ*wspFpelpsl* parent in the biofilm bead model, all SCVs with large genome deletions had reduced fitness in this assay (Fig S6B). Lastly, we used SCV 52 and SCV 57 as representative SCVs with large genomic deletions and assessed survival when exposed to host antimicrobials. Neither SCV showed differences in survival when treated with H_2_O_2_, compared to the Δ*wspFpelpsl* parent (Fig S6C). However, both SCVs showed increased survival when treated with serum (Fig S6C). In addition to the large genomic deletions, SCV 52 has a base pair insertion in *wzy*, while SCV 57 has a 3,332 base pair deletion, resulting in the deletion of *wbpH*, *wbpG, hisF2, hisH2, wzx* and *wzy* (Table 2). We therefore hypothesize that the increased tolerance to serum is due to the co-occurring *wzy* mutation, like what we observed for the *wzy*_620insC_ mutation (Fig 8), rather than due to contributions from the large genomic deletions.

Therefore, at this time from our *in vitro* assays, we are unable to determine the fitness benefits associated with these large genomic deletions.

### Evidence for filamentous phage activity

Lastly, we observed that three of the sequenced SCVs (SCVs 48, 49, and 50) displayed a significant increase in reads mapping to a specific region of the genome, with approximately 10- 20 fold increased coverage (Fig 9A). Furthermore, SCVs 69 and 70 displayed approximately a 4 fold increased coverage at the same genomic region. Review of this region indicated that the reads mapped to the genes encoding the Pf6 prophage, which was observed as increased coverage across the entire prophage region, compared to the flanking bacterial genes (Fig 9B).

**Figure 9:**
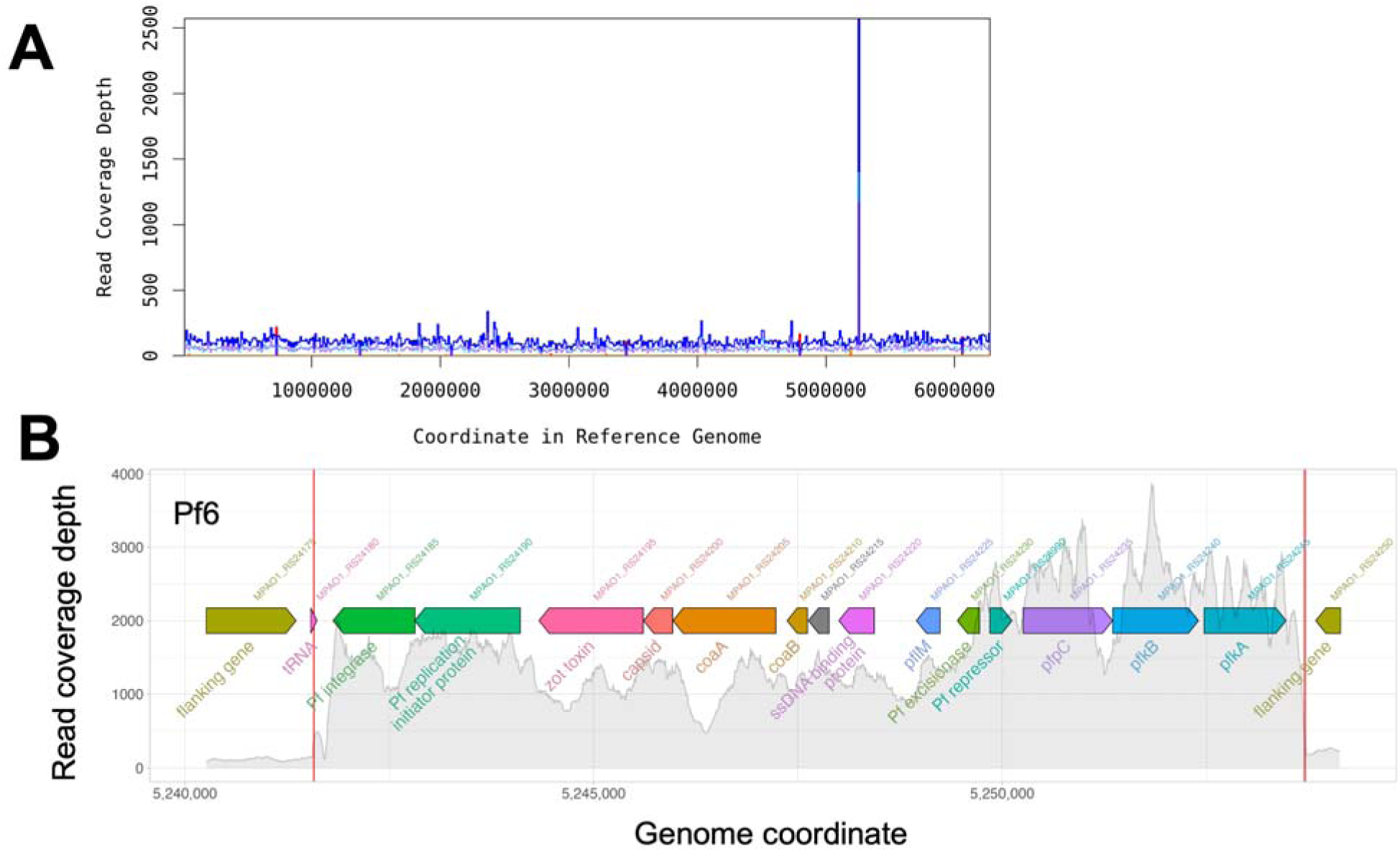
SCVs have increased read coverage across Pf6 encoded genes. Read coverage analysis of SCV 50. **(A)** Read coverage of SCV 50 sequences mapped to the reference MPAO1 genome. **(B)** Read coverage of SCV 50 sequences aligning to Pf6 genes in the reference genome. Colored arrows indicate the genes in this region, red lines indicate the genome regions corresponding to the encode prophage. Grey bars indicate the number of sequences aligning to the genome. Similar coverage was observed for SCVs 48, 49, 50. SCV 69 and 70 had coverage increases in this region but at lower levels. SCV 50 is depicted here as a representative of these isolates.

Notably, we did not observe increased coverage of Pf6 or Pf4 in the Δ*wspFpelpsl* parent strain. (Fig S7). Together these data suggest that Pf6 filamentous phage was replicating independently of the host bacterial genome. This was not inherent to the parental mutant background, but appeared to be activated during chronic infection of the porcine wound.

## Discussion

Here we demonstrate that in a porcine full-thickness thermal injury wound model, a triple gene mutant deficient in biofilm formation, Δ*wspFpelpsl,* undergoes adaptive evolution by acquiring mutations that alter the outer membrane; either hyper piliation or loss of O-antigen, that restores the deficient biofilm phenotype. For *pilU* mutants, this increased biofilm phenotype was associated with enhanced fitness compared to the parent, while for *wzy* mutants this was associated with increased tolerance to host antimicrobial products. To our knowledge, this is the first time these phenotypes and fitness benefits have been identified for *pilU* and *wzy* mutants, contributing to the growing understanding of *P. aeruginosa* adaptation to infection, and the genotype-phenotype associations that are under selection. This data adds to the growing evidence that the hyperbiofilm phenotype is adaptive in chronic infections and that *P. aeruginosa* has redundant pathways to generate this phenotype. Our data is also supportive of using variant colony morphology as a screen to identify these adaptive mutations from complex clinical samples (29).

We also observe a striking degree of mutational parallelism (Fig 5), at both the biosynthetic pathway (T4P and LPS; Fig 3) and gene (*pilU* and *wzy*; Fig 4) level, indicating the strong selective pressures experienced by these pathways in a chronic wound infection. Interestingly, similar mutational parallelism in *wzy* was identified in PAO1 in response to lytic phage predation (30). Previous studies identified mutations in other LPS biosynthetic genes and in T4P genes, also similar to what we observed here (30). Together these data suggest that mutations in T4P and LPS genes have pleiotropic phenotypes that are selected to expand the niche colonization and survival of *P. aeruginosa*.

Mutations in these pathways are also under selection in *P. aeruginosa* CF lung infections, where similar parallel evolution is observed (2, 3). However, interestingly this mutational parallelism is observed across more extensive genes, with previous studies identifying 52 genes that are repeatedly targeted during CF lung infections (3). By contrast, using the porcine wound model we identified mutational parallelism in 4 genes, *wspA* (18), *retS* (19), and here in *pilU* and *wzy*. This could be due to different selective pressures between a wound and lung infection environment, differences between adaptive evolution of lab strains compared to clinical isolates, or specific to our wound model. However, a caveat of our findings is that we were biased to those mutations that resulted in a variant colony morphology. Utilizing unbiased whole population metagenomic sequencing could reveal more complex evolutionary dynamics in the porcine wound model and will be the focus of future studies.

We also observed significant genome rearrangement of a subpopulation of SCVs during infection. Firstly, we observed large genome deletions, that occurred in similar regions of the genome (Table 2, Fig 5). Currently, we have been unable to determine the molecular mechanism for these deletions. However, this region of the *P. aeruginosa* genome has been identified as an extended nonessential region, that has been targeted for genome minimization and streamlining (31). Genome reduction of *P. aeruginosa* has been identified in CF lung infections as the isolates become host restricted (32). However, this was observed through the acquisition of pseudogenes, rather than genome deletions as we observed here. Furthermore, host restriction is typically observed over extended time frames of host colonization, with Armbruster *et al.* observing pseudogene evolution over a 30-month period (32). This suggests that the large genome deletions we observe here may not be a consequence of host restriction. Rather it seems to be reminiscent of genomic ‘black holes’ that were first described in *Shigella* spp (33). A large 190kb genome deletion was identified in *S. flexneria* and enteroinvasive *Escherichia coli* (EIEC) that was absent in non-pathogenic *E. coli*. This large deletion resulted in the removal of the *cadA* locus, conferring increased virulence of *S. flexneria* and EIEC strains (33). Genomic black holes have also been implicated in the evolution of pathogenic *Bacillus*, *Burkholderia, Bordetella, Rickettsiae, Mycobacterium* and *Chlamydia* spp by removing anti-virulence genes (34–36). Similarly, large genomic deletions in *P. aeruginosa* have been identified as a response to phage infection, where the deletion removes a gene, or gene cluster, essential for phage entry into the cell, resulting in phage resistance (37, 38). It is therefore interesting to speculate that the large genome deletions that we identified here confer increased persistence in the wound of the Δ*wspFpelpsl* parent, which is deficient in biofilm formation, by removal of anti-virulence, or anti-colonization genes. Deletion of these genes may also lead to cross protection to phage infection. This would also account for these deletions occurring in the same region of the genome, across different isolates.

Secondly, we observed evidence of Pf6 transitioning to the replicative form (Fig 9). Filamentous phage exist in the host as either two forms; a ssDNA infectious form that is integrated in the host genome as a prophage, or a dsDNA replicative form (RF) that is plasmid-like and replicates independently of the host genome (39). The transition between these forms is associated with changes in the environmental or growth conditions of the host bacterium (40, 41). This suggests that the wound environment induces the RF of Pf6 in Δ*wspFpelpsl*. Furthermore, our detection of the Pf6 RF from the wound is likely an under-representation, as this form can transition back to the prophage form within the host. The increased replication of Pf6 in Δ*wspFpelpsl* SCVs isolated from porcine wounds is notable, as filamentous phage produce phenotypes associated with increased biofilm formation, antibiotic resistance, resistance to phagocytosis and altered mammalian inflammatory responses (39, 42–44). Further, as T4P is the receptor for Pf phage, it is plausible that the T4P mutations observed here may have been selected as defenses against Pf superinfection, which is costly for fitness (45). Lastly, filamentous phage, specifically Pf4, has been associated with the SCV phenotype (44, 46). Interestingly, we identified a number of SCVs with mutations in MPAO1_RS28990, which is implicated as a lambda repressor gene (Table 2). Genes with similar annotated domains control lysogeny (47). Together this suggests that Pf6 may contribute to the SCV phenotype in the absence of LPS or T4P mutations.

For *P. aeruginosa*, SCV is used synonymously with RSCV; where these variants are typically associated with mutations leading to increased c-di-GMP and subsequent increased production of Pel and Psl polysaccharides (48–50). Here we describe SCVs that arise independently of polysaccharide overproduction. This indicates that the general SCV phenotype is indicative of infection adaptation, independent of genotype. Due to our findings, we propose that the designator of RSCV be used to describe those variants that arise from mutations resulting in increased levels of c-di-GMP, with a wrinkled or rugose colony morphology, to more clearly distinguish between polysaccharide dependent (RSCV) and independent (SCV) small colony variants, and the associated fitness benefits.

## Materials and Methods

### Bacterial strains and plasmids

Bacterial strains and plasmids used this study are detailed in Table S1. Mutant and deletion constructs were made using NEBuilder HiFi DNA assembly (NEB). Complementing constructs were made by ligation using T4 ligase (NEB) according to standard protocols. Mutant and deletion constructs were incorporated into the *P. aeruginosa* genome by two-step allelic recombination. Complementation constructs were introduced into *P. aeruginosa* by electroporation. Plasmid selection was maintained using 100 μg/mL ampicillin or 10 μg/mL gentamicin for *E. coli* and 300 μg/mL carbenicillin or 50 μg/mL gentamicin for *P. aeruginosa*.

### Porcine thermal-injury chronic wound model

Protocols were performed in accordance with OSU IACUC approval. Pigs were wounded and monitored as previously described (20). Briefly, two pigs were subjected to thermal injury by applying a 2 x 2-inch metal dice heated to 150°C for 25s to the back of the pig, to achieve six (three down each side) full-thickness thermal wounds. Three days post wounding, wounds were inoculated topically with 250 μL 10^8^ CFU/mL of PAO1Δ*wspF*Δ*pelA*Δ*pslBCD* (21). Three 8mm punch biopsies were sampled from two wounds from each pig at 7-, 14-, and 35-days post inoculation. Biopsies were homogenized in 1mL PBS and serially diluted to enumerate for CFU/g tissue and screen for colony morphology variants. Variants were passaged twice on Luria agar (LA), and once on *Pseudomonas* isolation agar (PIA) to confirm the variant morphology was stable. Confirmed variants were stored at -80°C.

To image the variant colony morphology, 1μL of an overnight culture of representative variants was spotted onto PIA and incubated overnight at 37°C. Colonies were imaged using a Lecia x stereoscope, fitted with x camera. Images were captured using x software and processed using FIJI (51).

To determine the fitness of the small-colony variants in the wound, the selection rate (*r*) was calculated according to equation 1 (52).

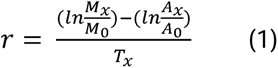

where *T* is time at day x, and *M* and *A* are the number of mutant and ancestor cells, respectively, at days 0 and x. As the number of mutants in the wound at day 0 is unknown, *s* was estimated using mutant to parent ratios of 1:10^5^, 1:10^6^, 1:10^7^ and 1:10^8^.

### Sequencing and analysis

DNA extraction and genome sequencing was performed as previously described (Gloag 2019). Briefly, genomic DNA was extracted using the DNeasy Blood and Tissue kit (Qiagen) following the manufacturer’s protocol. Isolate DNA was sequenced with the library preparation method according to Baym et al. (53) on an Illumina NextSeq 500. Paired-end 2x151 sequencing reads were quality filtered and trimmed with Trimmomatic v0.36 (settings: LEADING:20 TRAILING:20 SLIDINGWINDOW:4:20 MINLEN:70) (54) then variants were called with *breseq* v0.30.0 using default settings (55). We used GCF_016107485.1 as the MPAO1 reference assembly.

For Pf6 analysis, raw reads were trimmed using Trimmomatic v0.36 (settings: LEADING:3 TRAILING:3 SLIDINGWINDOW:4:15 MINLEN:36) (54) and reads were mapped using breseq v0.39.0 with default settings (55). We used MPAO1 (GCF_016107485.1) as the reference genome. Read depth was calculated by taking the bam file generated by breseq, and then averaging reads mapped across a 10 bp window using bedtools v2.26.0 (56) and samtools v1.21 (57).

### Twitching motility assay

Twitching motility was quantified as described previously (58). Briefly, a pipette tip was coated with bacterial culture grown overnight on LA, and stabbed through 1% LA plates (LB solidified with 1% agar), down to the interstitial space between the plastic and agar. Plates were incubated overnight at 37°C in a humidified chamber. The zone of twitching motility was observed as a halo at the interstitial space, and the diameter measured in mm. Three biological replicates were performed.

### LPS Western Blot

Overnight cultures were normalized to an OD_600nm_ 0.5 in LB and centrifuged at 10,600 x g for 10 minutes. Pellets were resuspended in 1X Laemmli buffer supplemented with β-mercaptoethanol, boiled at 100°C for 15 minutes, and then allowed to cool to room temperature for 15 minutes. Proteinase K (10mg/mL) was added and samples were incubated at 59°C for 1 hour. Samples were frozen at -20°C until further use.

Samples were electrophoresed using a 12% Mini-PROTEAN TGX gel (Biorad) for 1.25 hours at 120V. After electrophoresis, samples were transferred to a nitrocellulose membrane and the membrane was blocked for 1 hour in 5% nonfat milk at room temperature. Membranes were incubated in mouse anti-PA antigen O5 (My BioSource) overnight at 4°C. After primary antibody incubation, membranes were washed three times and incubated in goat anti-mouse secondary antibody conjugated to HRP for 1 hour at room temperature. Membranes were washed three times before chemiluminescent detection with Supersignal Pico (ThermoFisher) was completed. Images were acquired on a ChemiDoc Imaging System (BioRad).

### Biofilm quantification

Overnight cultures were normalized to an OD_600nm_ 0.5 in Jensen’s defined media (pH 7.3) (59). 100μL was transferred to a well of a 96-well plate and incubated for 24h at 37°C in a humidified chamber. Biofilms were stained with 120μL 0.1% crystal violent for 30min. Biofilms were washed three times in PBS and bound crystal violet dissolved in 150μL 100% ethanol for 30min. OD_590nm_ was measured on a SpectraMax i3 plate reader (Molecular Devices). Three biological replicates were performed, each with three technical replicates. Biofilm biomass was expressed as a percentage of PAO1, which was set to 100%.

### Biofilm competition

Parental strains were tagged with *lacZ* on the *attB* site using miniCTX::*lacZ* (60). The vector backbone was excised using pFLP2 (61).

Overnight cultures were normalized to an OD_600nm_ 1, and 50μL of competing strains were transferred to 5mL LB containing a 7mm polystyrene bead and incubated at 37°C in a rolling culture drum. After 24h incubation, biofilm-coated bead was transferred to 5mL LB with a second bead and incubated for another 24h (27). CFUs were enumerated at time 0 (inoculum culture) 24 and 48h on LA supplemented with 100μg/mL X-Gal. To quantify CFUs of biofilms, beads were sonicated in 1mL PBS in a water bath sonicator for 1min at 50% power. Biofilms were further disrupted by passing the sonicated culture through a 22” gauge needle. The selection coefficient (*s*) was calculated according to equation 1. Five biological replicates were performed, each with three technical replicates.

### Antimicrobial susceptibility

Overnight cultures were normalized to an OD_600nm_ 0.5 in LB. Samples were treated with 100% pig serum (Giboc) or 2.5% hydrogen peroxide for 1 hour at 37°C. PBS treatment was used as a control. Technical triplicates were serially diluted and enumerated for CFU/mL and expressed as percent survival relative to the PBS control. Three biological replicates were performed.

### Statistical analysis

Statistical analysis was performed using a one-way analysis of variance (ANOVA) with a Tukey’s multiple comparison post-hoc test, unless otherwise indicated in the figure legend. Analyses were performed using GraphPad Prism v.10 (GraphPad Software). Statistical significance was determined using a p-value < 0.05.

## Data availability

All sequencing data is available in NCBI SRA under BioProject number PRJNA1283160 and BioSample accession numbers SAMN49683420-SAMN49683459.

## Acknowledgments

ESG was funded by an AHA Career Development Award (19CDA34630005). DJW was funded by a NIH PHS grant R01AI186380.

## References

1. Kerr KG, Snelling AM. 2009. Pseudomonas aeruginosa: a formidable and ever-present adversary. J Hosp Infect 73:338–344.

2. Smith EE, Buckley DG, Wu Z. 2006. Genetic adaptation by Pseudomonas aeruginosa to the airways of cystic fibrosis patients. Proc Natl Acad Sci U S A.

3. Marvig RL, Sommer LM, Molin S, Johansen HK. 2015. Convergent evolution and adaptation of Pseudomonas aeruginosa within patients with cystic fibrosis. Nat Genet 47:57–64.

4. Sousa AM, Pereira MO. 2014. Pseudomonas aeruginosa Diversification during Infection Development in Cystic Fibrosis Lungs-A Review. Pathogens 3:680–703.

5. Randall TE, Eckartt K, Kakumanu S, Price-Whelan A, Dietrich LEP, Harrison JJ. 2022. Sensory Perception in Bacterial Cyclic Diguanylate Signal Transduction. J Bacteriol 204:e0043321.

6. Hickman JW, Tifrea DF, Harwood CS. 2005. A chemosensory system that regulates biofilm formation through modulation of cyclic diguanylate levels. Proc Natl Acad Sci U S A 102:14422–14427.

7. Kirisits MJ, Prost L, Starkey M, Parsek MR. 2005. Characterization of colony morphology variants isolated from Pseudomonas aeruginosa biofilms. Appl Environ Microbiol 71:4809– 4821.

8. Drenkard E, Ausubel FM. 2002. Pseudomonas biofilm formation and antibiotic resistance are linked to phenotypic variation. Nature 416:740–743.

9. O’Neal L, Baraquet C, Suo Z, Dreifus JE, Peng Y, Raivio TL, Wozniak DJ, Harwood CS, Parsek MR. 2022. The Wsp system of Pseudomonas aeruginosa links surface sensing and cell envelope stress. Proc Natl Acad Sci U S A 119:e2117633119.

10. Reinhardt A, Köhler T, Wood P, Rohner P, Dumas J-L, Ricou B, van Delden C. 2007. Development and persistence of antimicrobial resistance in Pseudomonas aeruginosa: a longitudinal observation in mechanically ventilated patients. Antimicrob Agents Chemother 51:1341–1350.

11. Ikeno T, Fukuda K, Ogawa M, Honda M, Tanabe T, Taniguchi H. 2007. Small and rough colony pseudomonas aeruginosa with elevated biofilm formation ability isolated in hospitalized patients. Microbiol Immunol 51:929–938.

12. Sheehan DJ, Janda JM, Bottone EJ. 1982. Pseudomonas aeruginosa: changes in antibiotic susceptibility, enzymatic activity, and antigenicity among colonial morphotypes. J Clin Microbiol 15:926–930.

13. Tielen P, Wibberg D, Blom J, Rosin N, Meyer A-K, Bunk B, Schobert M, Tüpker R, Schatschneider S, Rückert C, Albersmeier A, Goesmann A, Vorhölter F-J, Jahn D, Pühler A. 2014. Genome Sequence of the Small-Colony Variant Pseudomonas aeruginosa MH27, Isolated from a Chronic Urethral Catheter Infection. Genome Announc 2.

14. Tielen P, Narten M, Rosin N, Biegler I, Haddad I, Hogardt M, Neubauer R, Schobert M, Wiehlmann L, Jahn D. 2011. Genotypic and phenotypic characterization of Pseudomonas aeruginosa isolates from urinary tract infections. Int J Med Microbiol 301:282–292.

15. Brock MT, Fedderly GC, Borlee GI, Russell MM, Filipowska LK, Hyatt DR, Ferris RA, Borlee BR. 2017. Pseudomonas aeruginosa variants obtained from veterinary clinical samples reveal a role for cyclic di-GMP in biofilm formation and colony morphology. Microbiology 163:1613–1625.

16. Kasatvo AV, Gorovits ES, Kuznetsova MV, Timasheva OA, Sukhanov SG. 2015. Evaluation of biological properties of Pseudomonas aeruginosa strains isolated from patients with sternum and rib osteomyelitis. Zh Mikrobiol Epidemiol Immunobiol 69–74.

17. Boles BR, Thoendel M, Singh PK. 2004. Self-generated diversity produces “insurance effects” in biofilm communities. Proceedings of the National Academy of Sciences 101:16630–16635.

18. Gloag ES, Marshall CW, Snyder D, Lewin GR, Harris JS, Santos-Lopez A, Chaney SB, Whiteley M, Cooper VS, Wozniak DJ. 2019. Pseudomonas aeruginosa Interstrain Dynamics and Selection of Hyperbiofilm Mutants during a Chronic Infection. MBio 10.

19. Marshall CW, Gloag ES, Lim C, Wozniak DJ, Cooper VS. 2021. Rampant prophage movement among transient competitors drives rapid adaptation during infection. Sci Adv 7.

20. Roy S, Elgharably H, Sinha M, Ganesh K, Chaney S, Mann E, Miller C, Khanna S, Bergdall VK, Powell HM, Cook CH, Gordillo GM, Wozniak DJ, Sen CK. 2014. Mixed-species biofilm compromises wound healing by disrupting epidermal barrier function. J Pathol 233:331– 343.

21. Harrison JJ, Almblad H, Irie Y, Wolter DJ, Eggleston HC, Randall TE, Kitzman JO, Stackhouse B, Emerson JC, Mcnamara S, Larsen TJ, Shendure J, Hoffman LR, Wozniak DJ, Parsek MR. 2020. Elevated exopolysaccharide levels in Pseudomonas aeruginosa flagellar mutants have implications for biofilm growth and chronic infections. PLoS Genet 16:e1008848.

22. Pestrak MJ, Chaney SB, Eggleston HC, Dellos-Nolan S, Dixit S, Mathew-Steiner SS, Roy S, Parsek MR, Sen CK, Wozniak DJ. 2018. Pseudomonas aeruginosa rugose small-colony variants evade host clearance, are hyper-inflammatory, and persist in multiple host environments. PLoS Pathog 14:e1006842.

23. Whitchurch CB, Mattick JS. 1994. Characterization of a gene, pilU, required for twitching motility but not phage sensitivity in Pseudomonas aeruginosa. Mol Microbiol 13:1079–1091.

24. Islam ST, Taylor VL, Qi M, Lam JS. 2010. Membrane topology mapping of the O-antigen flippase (Wzx), polymerase (Wzy), and ligase (WaaL) from Pseudomonas aeruginosa PAO1 reveals novel domain architectures. MBio 1.

25. Han X, Kennan RM, Davies JK, Reddacliff LA, Dhungyel OP, Whittington RJ, Turnbull L, Whitchurch CB, Rood JI. 2008. Twitching motility is essential for virulence in Dichelobacter nodosus. J Bacteriol 190:3323–3335.

26. Chiang P, Habash M, Burrows LL. 2005. Disparate subcellular localization patterns of Pseudomonas aeruginosa Type IV pilus ATPases involved in twitching motility. J Bacteriol 187:829–839.

27. Poltak SR, Cooper VS. 2011. Ecological succession in long-term experimentally evolved biofilms produces synergistic communities. ISME J 5:369–378.

28. Whitchurch CB, Hobbs M, Livingston SP, Krishnapillai V, Mattick JS. 1991. Characterisation of a Pseudomonas aeruginosa twitching motility gene and evidence for a specialised protein export system widespread in eubacteria. Gene 101:33–44.

29. Branda SS, Vik S, Friedman L, Kolter R. 2005. Biofilms: the matrix revisited. Trends Microbiol 13:20–26.

30. Latino L, Midoux C, Hauck Y, Vergnaud G, Pourcel C. 2016. Pseudolysogeny and sequential mutations build multiresistance to virulent bacteriophages in Pseudomonas aeruginosa. Microbiology 162:748–763.

31. Csörgő B, León LM, Chau-Ly IJ, Vasquez-Rifo A, Berry JD, Mahendra C, Crawford ED, Lewis JD, Bondy-Denomy J. 2020. A compact Cascade-Cas3 system for targeted genome engineering. Nat Methods 17:1183–1190.

32. Armbruster CR, Marshall CW, Garber AI, Melvin JA, Zemke AC, Moore J, Zamora PF, Li K, Fritz IL, Manko CD, Weaver ML, Gaston JR, Morris A, Methé B, DePas WH, Lee SE, Cooper VS, Bomberger JM. 2021. Adaptation and genomic erosion in fragmented Pseudomonas aeruginosa populations in the sinuses of people with cystic fibrosis. Cell Rep 37:109829.

33. Maurelli AT, Fernández RE, Bloch CA, Rode CK, Fasano A. 1998. “Black holes” and bacterial pathogenicity: a large genomic deletion that enhances the virulence of Shigella spp. and enteroinvasive Escherichia coli. Proc Natl Acad Sci U S A 95:3943–3948.

34. Salimiyan RK, Farsiani H. 2022. Black holes”,“Genome fluidity”, and Evolution of Bacterial Species. Reviews in Clinical Medicine 9:146–155.

35. Day WA, Maurelli AT. 2014. Black holes and antivirulence genes: Selection for gene loss as part of the evolution of bacterial pathogens, p. 109–122. In Evolution of Microbial Pathogens. ASM Press, Washington, DC, USA.

36. Maurelli A. 2007. Black holes, antivirulence genes, and gene inactivation in the evolution of bacterial pathogens. FEMS Microbiol Lett 267:1–8.

37. Latino L, Essoh C, Blouin Y, Vu Thien H, Pourcel C. 2014. A novel Pseudomonas aeruginosa bacteriophage, Ab31, a chimera formed from temperate phage PAJU2 and P. putida lytic phage AF: characteristics and mechanism of bacterial resistance. PLoS One 9:e93777.

38. Le S, Yao X, Lu S, Tan Y, Rao X, Li M, Jin X, Wang J, Zhao Y, Wu N, Lux R, He X, Shi W, Hu F. 2014. Chromosomal DNA deletion confers phage resistance to Pseudomonas aeruginosa. Sci Rep 4.

39. Secor PR, Burgener EB, Kinnersley M, Jennings LK, Roman-Cruz V, Popescu M, Van Belleghem JD, Haddock N, Copeland C, Michaels LA, de Vries CR, Chen Q, Pourtois J, Wheeler TJ, Milla CE, Bollyky PL. 2020. Pf bacteriophage and their impact on Pseudomonas virulence, mammalian immunity, and chronic infections. Front Immunol 11:244.

40. Xue H, Xu Y, Boucher Y, Polz MF. 2012. High frequency of a novel filamentous phage, VCY φ, within an environmental Vibrio cholerae population. Appl Environ Microbiol 78:28– 33.

41. Jian H, Xu J, Xiao X, Wang F. 2012. Dynamic modulation of DNA replication and gene transcription in deep-sea filamentous phage SW1 in response to changes of host growth and temperature. PLoS One 7:e41578.

42. Secor PR, Michaels LA, Smigiel KS, Rohani MG, Jennings LK, Hisert KB, Arrigoni A, Braun KR, Birkland TP, Lai Y, Hallstrand TS, Bollyky PL, Singh PK, Parks WC. 2017. Filamentous bacteriophage produced by Pseudomonas aeruginosa alters the inflammatory response and promotes noninvasive infection*in vivo*. Infect Immun 85.

43. Burgener EB, Sweere JM, Bach MS, Secor PR, Haddock N, Jennings LK, Marvig RL, Johansen HK, Rossi E, Cao X, Tian L, Nedelec L, Molin S, Bollyky PL, Milla CE. 2019. Filamentous bacteriophages are associated with chronic Pseudomonas lung infections and antibiotic resistance in cystic fibrosis. Sci Transl Med 11.

44. Secor PR, Sweere JM, Michaels LA, Malkovskiy AV, Lazzareschi D, Katznelson E, Rajadas J, Birnbaum ME, Arrigoni A, Braun KR, Evanko SP, Stevens DA, Kaminsky W, Singh PK, Parks WC, Bollyky PL. 2015. Filamentous Bacteriophage Promote Biofilm Assembly and Function. Cell Host Microbe 18:549–559.

45. Kubota N, Scribner MR, Cooper VS. 2025. Filamentous cheater phages drive bacterial and phage populations to lower fitness. bioRxivorg.

46. Webb JS, Lau M, Kjelleberg S. 2004. Bacteriophage and phenotypic variation in Pseudomonas aeruginosa biofilm development. J Bacteriol 186:8066–8073.

47. Bell CE, Frescura P, Hochschild A, Lewis M. 2000. Crystal structure of the lambda repressor C-terminal domain provides a model for cooperative operator binding. Cell 101:801–811.

48. Häussler S. 2004. Biofilm formation by the small colony variant phenotype of Pseudomonas aeruginosa. Environ Microbiol 6:546–551.

49. Häußler S, Tümmler B, Weißbrodt H. 1999. Small-Colony Variants of Pseudomonas aeruginosa in Cystic Fibrosis. Clin Infect Dis.

50. Besse A, Groleau M-C, Trottier M, Vincent AT, Déziel E. 2022. Pseudomonas aeruginosa Strains from Both Clinical and Environmental Origins Readily Adopt a Stable Small-Colony- Variant Phenotype Resulting from Single Mutations in c-di-GMP Pathways. J Bacteriol 204:e0018522.

51. Schindelin J, Arganda-Carreras I, Frise E, Kaynig V, Longair M, Pietzsch T, Preibisch S, Rueden C, Saalfeld S, Schmid B, Tinevez J-Y, White DJ, Hartenstein V, Eliceiri K, Tomancak P, Cardona A. 2012. Fiji: an open-source platform for biological-image analysis. Nat Methods 9:676–682.

52. Cooper VS. 2018. Experimental Evolution as a High-Throughput Screen for Genetic Adaptations. mSphere 3.

53. Baym M, Kryazhimskiy S, Lieberman TD, Chung H, Desai MM, Kishony R. 2015. Inexpensive multiplexed library preparation for megabase-sized genomes. PLoS One 10:e0128036.

54. Bolger AM, Lohse M, Usadel B. 2014. Trimmomatic: a flexible trimmer for Illumina sequence data. Bioinformatics 30:2114–2120.

55. Deatherage DE, Barrick JE. 2014. Identification of mutations in laboratory-evolved microbes from next-generation sequencing data using breseq. Methods Mol Biol 1151:165– 188.

56. Quinlan AR, Hall IM. 2010. BEDTools: a flexible suite of utilities for comparing genomic features. Bioinformatics 26:841–842.

57. Li H, Handsaker B, Wysoker A, Fennell T, Ruan J, Homer N, Marth G, Abecasis G, Durbin R, 1000 Genome Project Data Processing Subgroup. 2009. The Sequence Alignment/Map format and SAMtools. Bioinformatics 25:2078–2079.

58. Semmler AB, Whitchurch CB, Mattick JS. 1999. A re-examination of twitching motility in Pseudomonas aeruginosa. Microbiology 145 (Pt 10):2863–2873.

59. Jensen SE, Fecycz IT, Campbell JN. 1980. Nutritional factors controlling exocellular protease production by Pseudomonas aeruginosa. J Bacteriol 144:844–847.

60. Hoang TT, Kutchma AJ, Becher A, Schweizer HP. 2000. Integration-proficient plasmids for Pseudomonas aeruginosa: site-specific integration and use for engineering of reporter and expression strains. Plasmid 43:59–72.

61. Hoang TT, Karkhoff-Schweizer RR, Kutchma AJ, Schweizer HP. 1998. A broad-host-range Flp-FRT recombination system for site-specific excision of chromosomally-located DNA sequences: application for isolation of unmarked Pseudomonas aeruginosa mutants. Gene 212:77–86.

