## Supplemental table and figures for "*Pseudomonas aeruginosa* biofilm-deficient mutants undergo parallel adaptation during chronic infection"

**Supplemental Table 1: List of strains and plasmids**

| **Strains and plasmids** | **Relevant genotype and/or characteristic** | **Reference** |
| --- | --- | --- |
| ***P. aeruginosa* strains** |  |  |
| MPAO1 | Colin Mannoil PAO1 strain. Referred to in the text as PAO1 |  |
| PAO1∆*pelA*∆*pslBCD* | Markerless deletion of *pelA* and *pslBCD.* Referred to in the text as ∆*pel*∆*psl* | (1) |
| PAO1∆*wspF* | Markerless deletion of *wspF* | (2) |
| PAO1∆*wspF*∆*pelA*∆*pslBCD* | Markerless deletion of *wspF, pelA* and *pslBCD.* Referred to in the text as ∆*wspF*∆*pel*∆*psl* | (2) |
| PAO1 *pilU*_∆560-570_ | 11bp deletion in *pilU*; 560-570/1149 | This study |
| PAO1 ∆*pelA*∆*pslBCD* *pilU*_∆560-570_ | Markerless deletion of *pelA* and *pslBCD*. 11bp deletion in *pilU*; 560-570/1149 | This study |
| PAO1 ∆*wspF pilU*_∆560-570_ | Markerless deletion of *wspF*. 11bp deletion in *pilU*; 560-570/1149 | This study |
| PAO1 ∆*wspF*∆*pelA*∆*pslBCD* *pilU*_∆560-570_ | Markerless deletion of *wspF*, *pelA* and *pslBCD*. 11bp deletion in *pilU*; 560-570/1149 | This study |
| PAO1∆*pilU* | Markerless deletion of *pilU* | This study |
| PAO1∆*pilT* | Markerless deletion of *pilT* | This study |
| PAO1 *wzy*_620insC_ | C bp insertion in *wzy;* (C)6→7 620/1317 | This study |
| PAO1 ∆*pelA*∆*pslBCD wzy*_620insC_ | Markerless deletion of *pelA* and *pslBCD*. C bp insertion in *wzy*; (C)6→7 620/1317 | This study |
| PAO1 ∆*wspF wzy*_620insC_ | Markerless deletion of *wspF*. C bp insertion in *wzy*; (C)6→7 620/1317 | This study |
| PAO1 ∆*wspF*∆*pelA*∆*pslBCD wzy*_620insC_ | Markerless deletion of *wspF*, *pelA* and *pslBCD*. C bp insertion in *wzy*; (C)6→7 620/1317 | This study |
| PAO1 *attB*::*lacZ* | *lacZ* inserted in the *attB* site | This study |
| PAO1∆*pelA*∆*pslBCD attB::lacZ* | Markerless deletion of *pelA* and *pslBCD*. *lacZ* inserted in the *attB* site | This study |
| PAO1∆*wspF* *attB::lacZ* | Markerless deletion of *wspF*. *lacZ* inserted in the *attB* site | This study |
| PAO1∆*wspF*∆*pelA*∆*pslBCD* *attB::lacZ* | Markerless deletion of *wspF*, *pelA* and *pslBCD*. *lacZ* inserted in the *attB* site | This study |
| ***E. coli* strains** |  |  |
| NEB5𝛼 | Cloning strain | NEB |
| S17 | Host for conjugation |  |
| **Plasmids** |  |  |
| pEX18Ap | Allelic exchange vector with *sacB*, and *bla* | (3) |
| pEX18Ap::*pilU*_∆560-570_ | Construct to re-create the *pilU* mutation. 500bp upstream and downstream the deletion site was cloned into pEX18Ap as a EcoRI-HindIII fragment | This study |
| pEX18Ap::∆*pilU* | Deletion construct for *pilU*. 1000bp upstream and downstream of *pilU* were cloned into pEX18Ap as a EcoRI-HindIII fragment | This study |
| pEX18Ap::∆*pilT* | Deletion construct for *pilT*. 1000bp upstream and downstream of *pilT* were cloned into pEX18Ap as a EcoRI-HindIII fragment | This study |
| pEX18Gm | Allelic exchange vector with *sacB*, and *aacC1* | (3) |
| pEX18Gm::*wzy*_620insC_ | Construct to re-create the wzy mutation. Mutant allele of *wzy* was amplified from SCV 79 and cloned in pEX18Gm | This study |
| pHERD20T::*mRuby3* | Empty vector control | Lori Burrows |
| pHERD20T::*mRuby3-pilU* | Complementing construct for *pilU* | Lori Burrows |
| miniCTX::*lacZ* | Construct to label strains with *lacZ*, *tet* | (4) |
| pUCP18 | Broad host range cloning vector, *Ap* | (5) |
| pUCP18::*wzy* | Complementing *wzy* construct. *wzy* was cloned into pUCP18 as a EcoRI-XbaI fragment | This study |

**Supplemental figures**

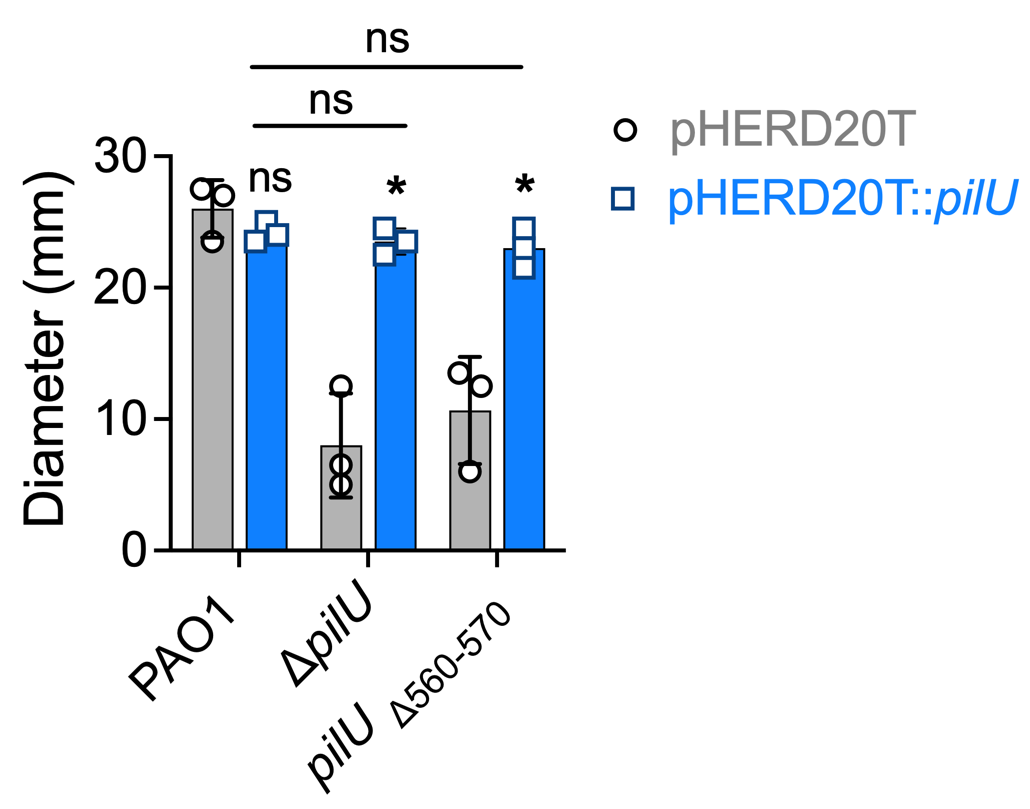

**Figure S1: Complementation of *pilU* mutants.** Diameter of the interstitial twitching zone of strains containing either the empty vector (pHERD20T) or the vector containing a wild type copy of *pilU* (pHERDT20T::*pilU*). Data presented as the mean ± SD. Individual points indicate the mean of each biological replicate. N=3. Significance was determined using a two-way ANOVA. * indicates *p* < 0.05, ns indicates no significance. Comparisons are to the empty vector control, unless indicated otherwise on the graph.

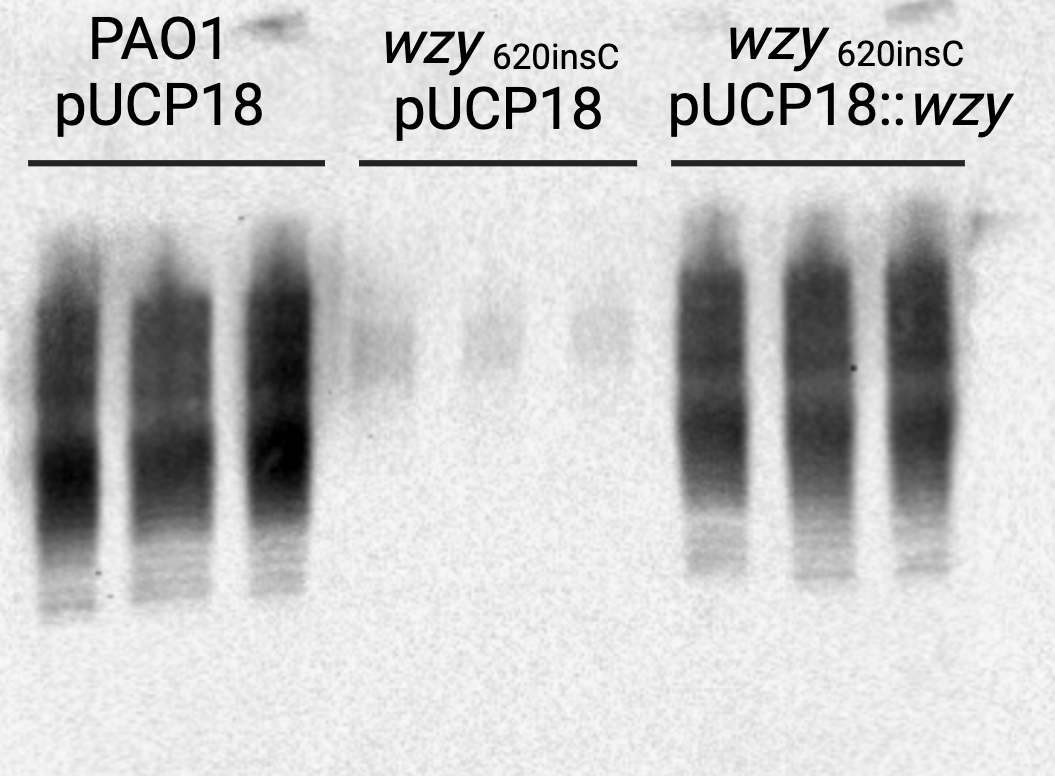

**Figure S2: Complementation of *wzy* mutation.** Western blot detecting O-antigen production in strains containing either the empty vector control (pUCP18), or the vector containing a wild type copy of *wzy* (pUCP18::*wzy*). Three lanes per strain indicate biological replicates. N=3.

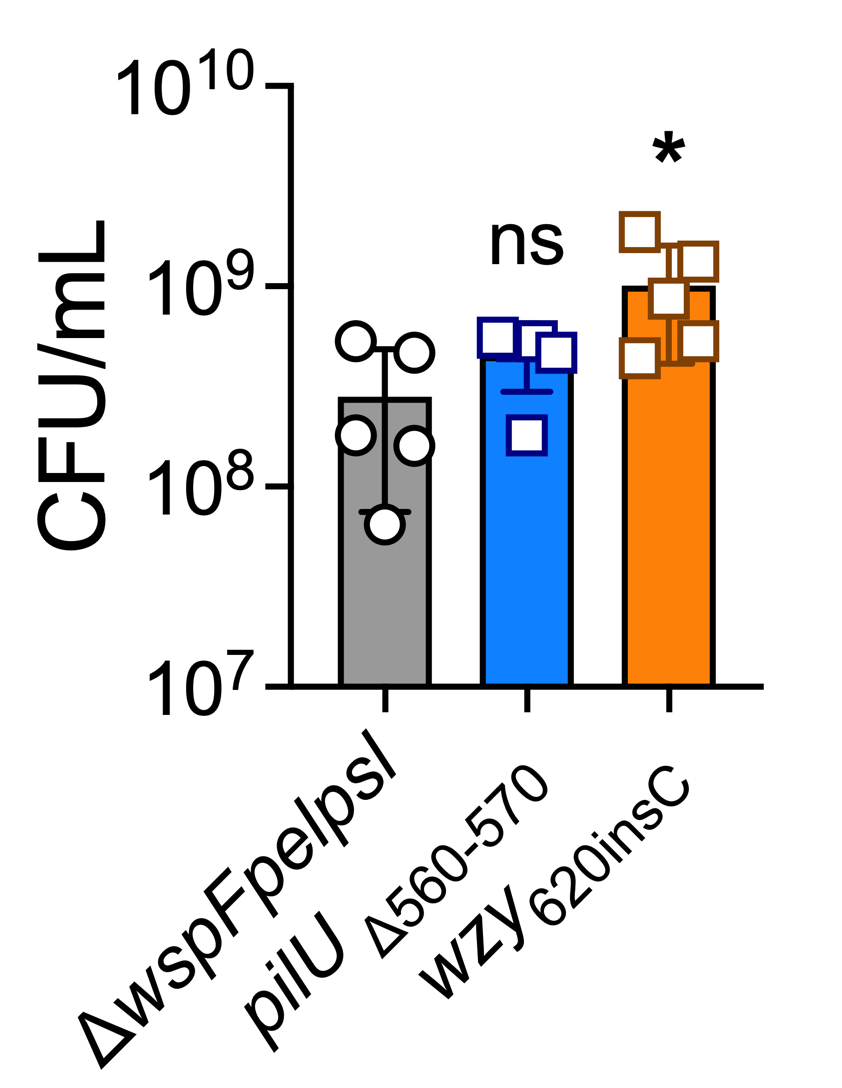

**Figure S3: Quantification of biofilm bacteria.** Biofilms were grown in a 96-well plate for 24h in Jensen’s media. Biofilm biomass was quantified by enumerating for CFU/mL. Data presented as mean ± SD. Individual data points indicates biological replicates, which is the mean of triplicate technical replicates. N = 5. * *p*-value <0.05, ns indicates no significance, compared to Δ*wpsFpelpsl*.

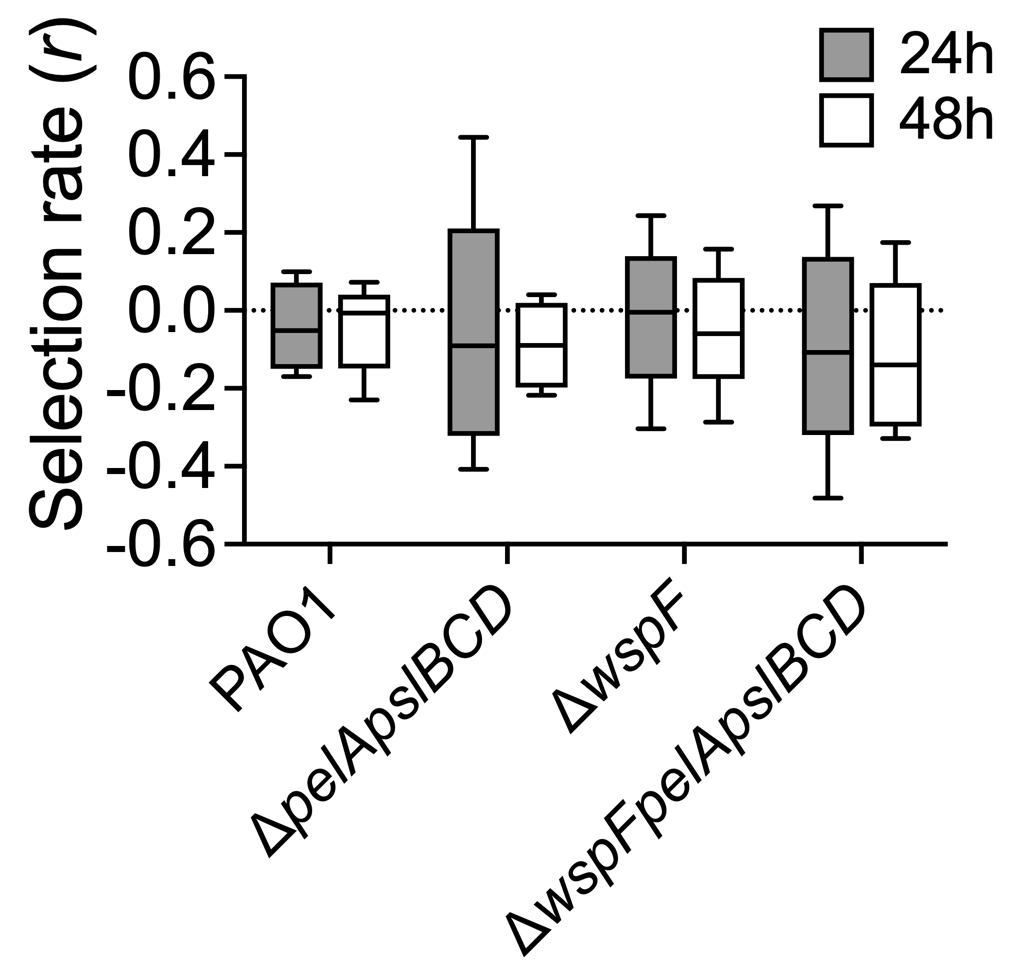

**Figure S4: *lacZ* tag does not affect fitness of the parent strain.** Selection rate (*r*) of the parent *P. aeruginosa* strains, competed pairwise against the *lacZ* tagged counterpart in a biofilm for 24 and 48h. N = 5.

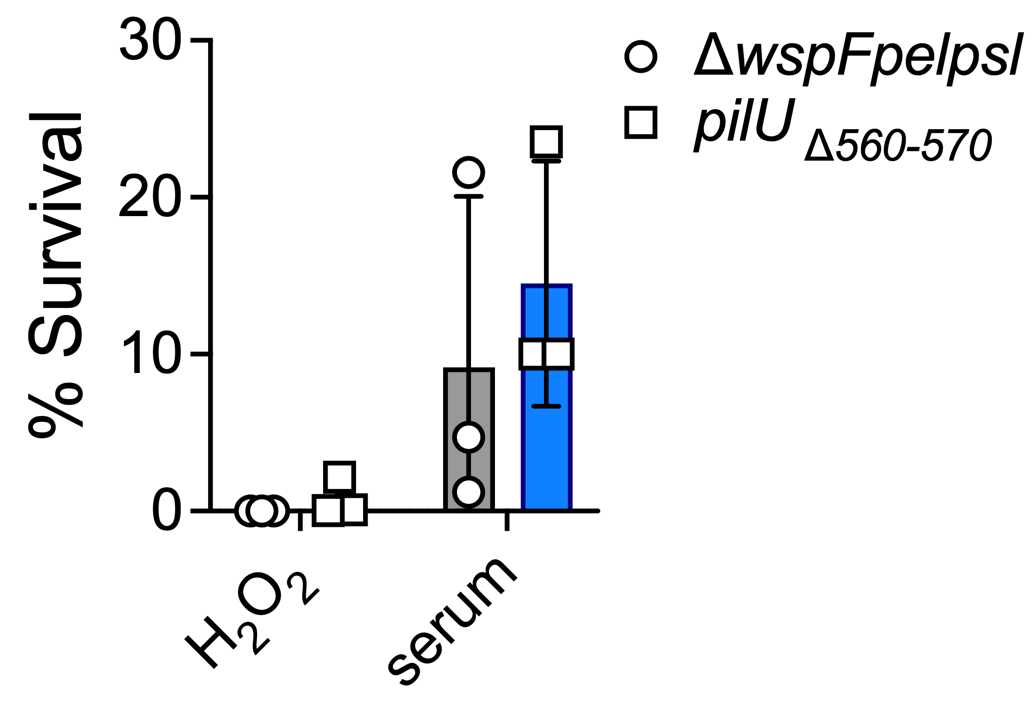

**Figure S5: *pilU*_Δ560-570_** **mutation is not protective against host antimicrobial products.** Δ*wspFpelpsl* and Δ*wspFpelpsl* *pilU*_Δ560-570_ was treated with either 2.5% H_2_O_2_ or 100% serum for 1h. Bacterial viability was enumerated by CFU/mL and expressed as percent survival relative to PBS untreated control. N = 3. Data presented as mean ± SD. Individual data points indicates biological replicates, which are the average of three technical replicates.

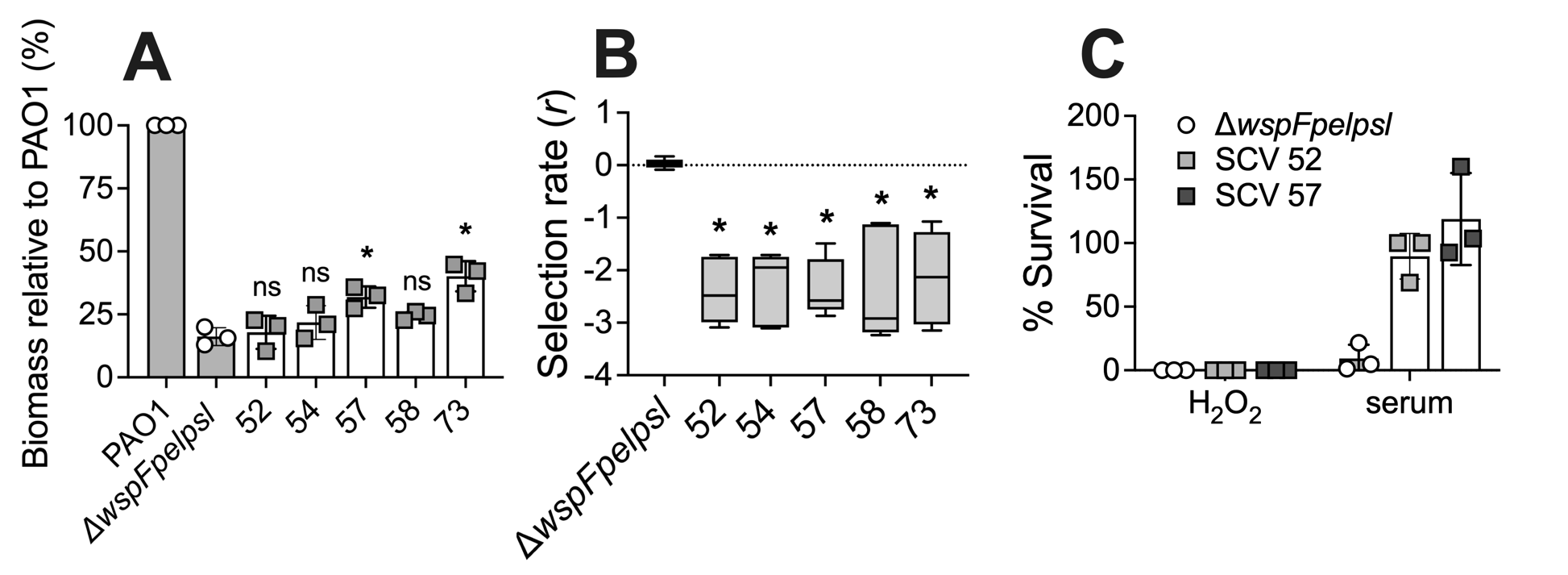

**Figure S6: Large genome deletions do not confer increased fitness *in vitro*. (A)** Biofilms were grown in a 96-well plate for 24h in Jensen’s media. Biofilm biomass was quantified by crystal violet staining. Biomass expressed as a percentage relative to PAO1. Data presented as mean ± SD. Individual data points indicates biological replicates, which are the average of four technical replicates. N = 3. * *p*-value <0.05, ns indicates no significance, compared to the parent background. **(B)** Selection rate (*r*) of indicated SCVs, competed pairwise against the parent in a biofilm for 48h. N = 5. * *p*-value <0.05, compared to the Δ*wspFpelpsl* competition. **(C)** Δ*wspFpelpsl* and representative SCVs were treated with either 2.5% H_2_O_2_ or 100% serum for 1h. Bacterial viability was enumerated by CFU/mL and expressed as percent survival relative to PBS untreated control. N = 3. Data presented as mean ± SD. Individual data points indicates biological replicates, which are the average of three technical replicates.

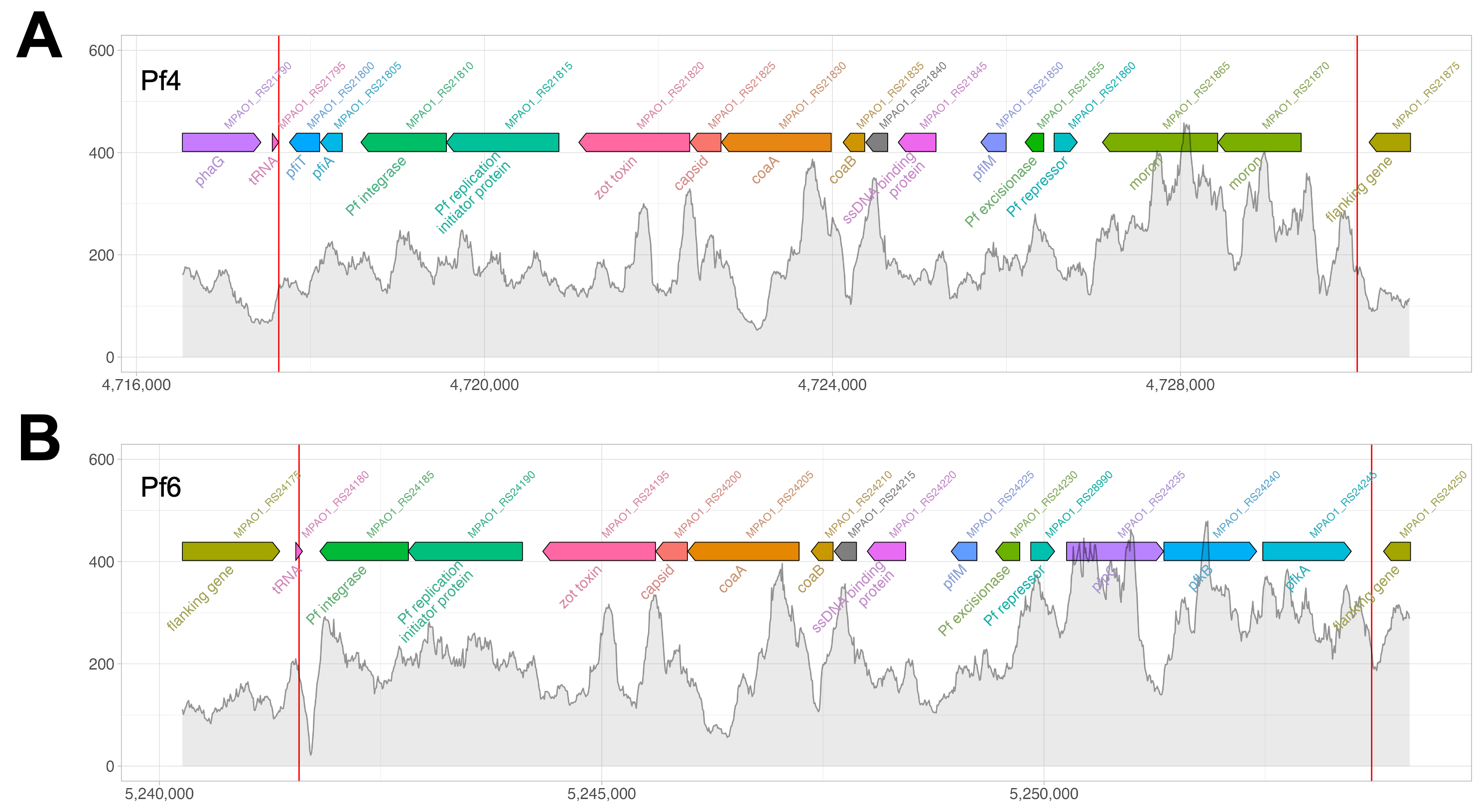

**Figure S7: Δ*wspFpelpsl* does not have increased read coverage across Pf4 or Pf6 encoded genes.** Read coverage of Δ*wspFpelpsl* sequences aligning to **(A)** Pf4 and **(B)** Pf6 genes in the reference genome. Colored arrows indicate the genes in this region, red lines indicate the genome regions corresponding to the encode prophage. Grey bars indicates the number of sequences aligning to the genome.

**References**

1.	Mishra M, Byrd MS, Sergeant S, Azad AK, Parsek MR, McPhail L, Schlesinger LS, Wozniak DJ. 2012. Pseudomonas aeruginosa Psl polysaccharide reduces neutrophil phagocytosis and the oxidative response by limiting complement-mediated opsonization. Cell Microbiol 14:95–106.

2.	Harrison JJ, Almblad H, Irie Y, Wolter DJ, Eggleston HC, Randall TE, Kitzman JO, Stackhouse B, Emerson JC, Mcnamara S, Larsen TJ, Shendure J, Hoffman LR, Wozniak DJ, Parsek MR. 2020. Elevated exopolysaccharide levels in Pseudomonas aeruginosa flagellar mutants have implications for biofilm growth and chronic infections. PLoS Genet 16:e1008848.

3.	Hoang TT, Karkhoff-Schweizer RR, Kutchma AJ, Schweizer HP. 1998. A broad-host-range Flp-FRT recombination system for site-specific excision of chromosomally-located DNA sequences: application for isolation of unmarked Pseudomonas aeruginosa mutants. Gene 212:77–86.

4.	Hoang TT, Kutchma AJ, Becher A, Schweizer HP. 2000. Integration-proficient plasmids for Pseudomonas aeruginosa: site-specific integration and use for engineering of reporter and expression strains. Plasmid 43:59–72.

5.	Schweizer HP. 1991. Escherichia-Pseudomonas shuttle vectors derived from pUC18/19. Gene 97:109–121.
